# Hippocampal TRPV1 channels in the modulation of contextual fear conditioning

**DOI:** 10.1101/2021.05.23.445340

**Authors:** Lia P. Iglesias, Heliana B. Fernandes, Aline S. de Miranda, Carlos A. Sorgi, Fabrício A. Moreira

**Author notes:** Correspondence: F.A. Moreira, Department of Pharmacology, Institute of Biological Sciences, Universidade Federal de Minas Gerais; Av. Pres. Antônio Carlos 6627, 31270-901 Belo Horizonte, MG, Brazil.

## Abstract

Psychiatric disorders have been linked to impairments in fear memory circuitry. Thus, pharmacological approaches that impair aversive memories have been investigated as new treatments. The TRPV1 channel modulates biological processes related to memory consolidation and retrieval. However, TRPV1 seems involved in memories generated by high intense conditioning. Anandamide (AEA), the main endocannabinoid, is an agonist of both, TRPV1 channels and CB1 receptors which are colocalized in several brain structures. Remarkably, **AEA has twenty-times more affinity for CB1 than for TRPV1, which may be involved in the intensity-dependent recruitment of this channel**. In order to evaluate the role of intensity of the conditioning in the recruitment of TRPV1, the animals were submitted to the contextual fear conditioning (CFC) and conditioned with low, moderate or high intensity. Before the retrieval a TRPV1 blocker was administered into the dorsal hippocampus (dHPC). The levels of AEA were quantified by Mass Spectrometry. The RNA levels of Arc, Zif and Trkb, involved in memory and plasticity, were quantified by PCR. Our results showed that TRPV1 blockers impair the retrieval of memory in animals conditioned with **moderate and high intensity** but not **low ones**. As revealed by Mass Spectrometry, this different recruitment among intensities seems to be associated with the levels of AEA released. Moreover, the impairment in freezing induced by blocking TRPV1 was prevented by a subeffective dose of the cannabinoid receptor CB1 antagonist which suggest that TRPV1 blockers act increasing AEA availability in the synaptic cleft to act through CB1 receptors. Despite blocking TRPV1 channels impairs freezing in **moderate and high intensities,** it increases the RNA levels of Arc, Zif and Trkb only in animals conditioned **with the moderate intensity**. In accordance, the treatment impairs **retrieval** in both intensities but only in **the moderate intensity** is able to prevent **the reinstatement**. Summarizing, our results suggested that intensity of the conditioning modulates AEA levels which in turns determines if TRPV1 will be recruited at the retrieval and which molecular pathways will be engaged due to TRPV1 blocking.

## INTRODUCTION

Post-traumatic stress disorder (PTSD) results from exposure to a severe traumatic event. This disorder is characterized by re-experiencing the trauma due to contextual reminders or intrusive thoughts as well as by psychophysiological reactivity and avoidance of trauma-related cues (Mahan & Ressler, 2012). This dysregulation in fear circuitry is related to higher rates of memory generalization (Lis et al., 2020) and memory persistence combined with impairments in extinction (Blechert et al., 2007; Milad et al., 2008). However, the role of intensity in promoting maladaptive fear memories or insufficient response to treatment is poorly understood.

The persistence of fear memory correlates with the intensity of the traumatic-like event (Bradley et al., 1992; Dos Santos Corrêa et al., 2019; Pereira et al., 2019; Ponce-Lina et al., 2020). In preclinical models, exposure to high-intense aversive stimuli is associated with faster generalization (Dos Santos Corrêa et al., 2019; Pedraza et al., 2016) and poor extinction performance (González-Franco et al., 2017). These effects are sustained by divergent recruitment of anatomical, cellular or molecular paths. For instance, different involvement of the amygdala (AMG), pre-frontal cortex (PFC) and hippocampus (HPC) was found under different intensities (Canto-de-Souza & Mattioli, 2016; Garín-Aguilar et al., 2014; Matos et al., 2019). In addition, corticosterone levels correlate with the intensity of conditioning (Cordero et al., 1998; Dos Santos Corrêa et al., 2019; Ponce-Lina et al., 2020), which in turns determines the ability of glucocorticoid antagonists to regulate fear memory (Cordero & Sandi, 1998). Similarly, intensity also modifies the modulation of fear memory by other drugs (Cruz-Morales et al., 1992; Quirarte et al., 1993), including antidepressants (Santos et al., 2006).

More recently, the transient receptor potential vanilloid type-1 channel (TRPV1) has also been implicated in the modulation of aversive responses (Iglesias et al., 2020; Moreira et al., 2012). The TRPV1 channel, a non-selective ionic channel with high permeability to Ca^2+^ (Caterina et al., 1997), is also included in the extended endocannabinoid system since it is activated by AEA, a component of the endocannabinoid system (ECS) (Zygmunt et al., 1999). The ECS includes the CB_1_ receptor (Matsuda et al., 1990) and the cannabinoid receptor CB_2_ (Munro et al., 1993), both of them Gi-protein-coupled receptors (Gonzalez, S., Sagredo, O., Gómez, M., Ramos, 2002), their endogenous ligands AEA (Devane et al., 1992) and 2-arachidonoylglycerol (2-AG) (Mechoulam et al., 1995), and the enzymes responsible for their metabolism, fatty acid amide hydrolase (FAAH) (Deutsch & Chin, 1993) and monoacylglycerol lipase (MAGL) (Tornqvist & Belfrage, 1976).

One of the main characteristics of this system is that endocannabinoids (eCB) are synthesized and released on demand by postsynaptic neurons (Pertwee, 2004), acting as retrograde messengers in their receptors located in presynaptic positions (Uchigashima et al., 2007). Moreover, TRPV1 channels and CB1 receptors are usually co-localized, the first in postsynaptic neurons (Toth et al., 2005), the second mainly in presynaptic positions (Cristino et al., 2006; Kauer & Gibson, 2009). Furthermore, CB1 and TRPV1 are activated by the same compound, AEA. Hence, drugs that interact with one of them can induce the recruitment of the other increasing the availability of AEA in the synaptic cleft. In this vein, the modulation of the ECS requires to consider I) CB1 and TRPV1 have usually opposite effects, II) AEA has twenty times more affinity for TRPV1 than for CB1, thus, the more the AEA released, the higher the probability to recruit TRPV1. Moreover, several alterations on the ECS were described in PTSD patients like altered levels of AEA (Neumeister et al., 2013), or the negative correlation between eCB levels and intrusive symptoms (Hill et al., 2013). These observations, among others, lead to propose this system as an interesting target to develop new treatments (Black et al., 2019; Chadwick et al., 2020; Lisboa et al., 2019).

For instance, Gobira et al. (2017) showed that AA-5-HT, a dual blocker of TRPV1 and FAAH, impaired fear expression, this was prevented by AM251, a CB1 antagonist and mimicked by the coadministration of subeffective doses of a FAAH blocker and a TRPV1 blocker (Gobira et al., 2017). Suggesting some involvement of TRPV1 channels in fear memory retrieval. However, evidences indicated that TRPV1 channels are recruited only after high intense conditioning (Genro et al., 2012; Terzian et al., 2009). In this sense, CB1 knock out presents an enhancement of fear memory related responses associated to high but not low intensities of training (Jacob et al., 2012), probably due to the recruitment of TRPV1 only in this intensity. Since higher levels of AEA are needed to recruit TRPV1 channels in the presence of CB1, this intensity-dependent recruitment may be associated to the levels of AEA released in each intensity. In this vein, it was reported that AEA levels seem higher after high intense conditioning (Morena et al., 2014). Despite the endocannabinoid system was extensively studied in relation to fear memory (Lutz et al., 2015), remains unknown if intensity of the traumatic-like event may modulate its signalling. Since intensity seems a key factor in maladaptive memory (Dos Santos Corrêa et al., 2019; González-Franco et al., 2017; Pedraza et al., 2016) and it may determine the effectivity of the treatment (Cordero & Sandi, 1998; Cruz-Morales et al., 1992; Quirarte et al., 1993; Santos et al., 2006). We tested the hypothesis that TRPV1 channels in the dHPC modulate contextual fear memory in an intensity-dependent manner. We also investigated how TRPV1 interacts with the ECS as well as the recruitment of inflammatory and neurotrophic factors.

## MATERIAL AND METHODS

(see Supplementary Material for more information)

### Animals

The experimental animals were 9 weeks-old male mice from C57B/L6J strain. They were located in plastic cages and kept in a controlled temperature of 24±2 C, with a dark-light cycle of 12/12h and free access to water and food. All the behavioural tests were performed during the light phase. The protocols were approved by the local ethics committee under the protocol number CEUA 176/2020.

### Drugs

The doses of TRPV1 blocker SB366791 (SB) were 1, 3 and 10 nmol. The sub-effective dose (75pmol) of AM251, a CB_1_ antagonist, was based on previous reports (Guimara et al., 2012; Hartmann et al., 2019). Both drugs were administered bilaterally into the dHPC and diluted in ethanol, chromophore, saline (1:1:18).

### Contextual Fear conditioning

The protocol was based on a previous report (Gobira et al., 2017). Briefly, in the first day was performed the conditioning, the animals were placed in the chamber and submitted to one of the protocols described in Table 1 (supplementary materials). Twenty-four hours later was performed the test. The animal was placed into the chamber during 5min without any intervention and recorded. In some experiments, a second test was performed 24h after the first.

The experiments involving the reinstatement were based on previous reports (Hitora-imamura et al., 2015; Vouimba & Maroun, 2011). One week after the test the mice were submitted to an extinction session, where they were placed in the chamber during 20 min. Twenty-for hours later was performed the reinstatement session where the animals return to the chamber and, after 3min, received one 500mA shock for 1s, after that the animals were kept in the chamber for one more minute. Twenty-four hours after the reinstatement session the animals were re-tested during 5min.

### Stereotaxic surgery and hippocampal administrations

The surgery and administration was performed as previously described (Gobira et al., 2017). The animals were anesthetized with ketamine and xylazine (i.p., 100mg/kg, 10mg/kg respectively). They were positioned in the stereotaxic apparatus; the skullcap was exposed and two guide cannulas were implanted. The stereotaxic coordinates were: AP −1.9mm, ML +1.5mm, DV −1.3mm (Paxinos & Franklin, 2003). A mixture of resin and acrylic was used to cover the skullcap. The animals were allowed to recover for 7 days before any other manipulation.

In order to perform the bilateral administration, the animal was immobilized and two dental needles were located into the cannulas. A volume of 0.25μl was administered using Hammilton microsyringes connected to an infusion pump in a flow of 0.25μl/min. The injector needles were retired 30s after the administration was concluded to avoid reflux.

### Immunofluorescence

The protocol was based on Fogaça et al. (Fogaça et al., 2012). Briefly, animals were prefunded with 100ml of phosphate buffered saline (PBS) and 50ml of paraformaldehyde (PFA) 4%. Coronal sections of 50 μm obtained in a vibratome were kept in PFA 2% at 4°C. The sections, were submitted to antigen retrieval, permeabilization, glycine 0.01M blocking and BSA5% blocking. The sections were incubated in primary antibodies diluted in BSA5% blocking solution for 72h: VR1 in goat (Santa Cruz, 1:30) + CB1 in rabbit (Invitrogen, 1:1000) or VR1 in goat (Santa Cruz, 1:30) + NeuN in mouse (Invitrogen, 1:500). After 72h of incubation the sections were incubated with secondary antibodies diluted in blocking solution for two hours: Alexa 594 anti-rabbit (Invitrogen, 1:1000), Alexa 488 High Cross Absorbed (Invitrogen, 1:1000) and Alexa 594 anti-mouse (Invitrogen, 1:1000). Finally, the sections were mounted using FlouroumntG.

### PCR

The lyse and homogenization of the samples was performed in accordance with TRIzol Reagent User Guide and the samples were stored at −80°C. M-MLV reverse transcriptase kit (Invitrogen) was used to obtain the cDNA. Quantitative PCR was performed using the iTaq™ Universal SYBR® Green Supermix (BioRad) in the CFX96 Touch™ Real Time detection system (BioRad). The data was analysed as previously described by Livak & Schmittgen (2001), the relative levels of RNA were calculated considering Rpl32.

### ELISA

The samples were homogenised and the supernatant was collected and stored at −80°C. The plate was first sensibilize. After that, the plate was blocked with blocking solution for 1h at RT. The samples were diluted 1:3 and 50 μl/well was added. The plate was kept at 4°C o.n followed by an incubation with streptavidin. Finally, it was incubated with 0.3 μg/ml of OPD diluted in citrate buffer and H2O2 (5:1). The reaction was interrupted with stop solution, H2SO4 1M. The plate was read at 490nm. The final concentrations were obtained using 4-parameter analysis.

### High performance liquid chromatography and Mass Spectrometry (HPLC-MS)

As previously described by de Oliveira et al., (2020). The samples were homogenized in H_2_O and MeOH and purified. After the samples were dried, they were resuspended in 100μl of MeOH and injected into the HLPC followed by MS. The mobile phases were water and acetonitrile containing 0.1% of formic acid. The MS was operated in positive mode.

### Statistical Analysis

The results were analysed by the student’s t-test or by analysis of variance (ANOVA) followed by Bonferroni post-hoc test, as appropriate. Correlation analysis were performed using Pearson’s. The statistical significance was set at p<0.05. The data are presented as mean and s.e.m.

## RESULTS

### Intensity-dependent recruitment of TRPV1 channels in the dHPC

Previous results from our group showed that AA-5HT, a dual blocker of TRPV1 and FAAH, administered into the dHPC impairs the retrieval of fear memory in the CFC (Gobira et al., 2017). In order to investigate if blocking TRPV1 without increasing AEA is sufficient to induce this impairment, we replicate the experiment describe by Gobira et al. (2017) (low intensity) using as treatment SB, a selective TRPV1 blocker. Our results revealed that blocking TRPV1 channels in the dHPC without increasing AEA levels did not induce alterations in freezing behaviour (Fig.1B) [F(3,37)=0.8317, p=0.4850].

**Figure 1.**
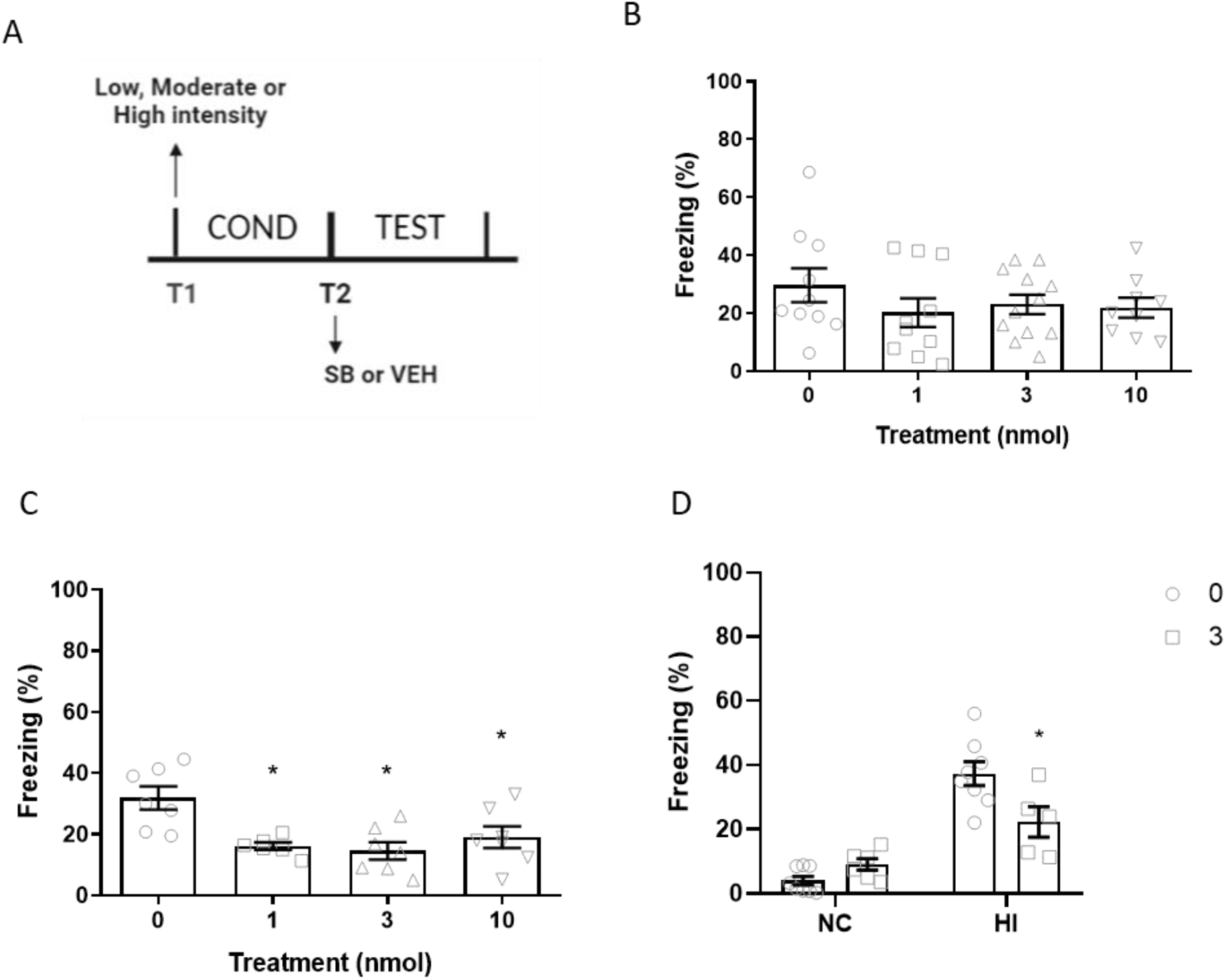
Intensity dependent recruitment of TRPV1 channels in the retrieval of contextual fear memory. Animals were conditioned with low (LI), moderate (MI), high intensity (HI) or not-conditioned (NC), SB was administered into the dHPC before retrieval. **A)** Experimental design **B)** Freezing levels of animals conditioned with low intensity [F(3,22)=1.161, p=0.3468],], n=6-7. **C)** Freezing levels of animals conditioned with moderate intensity [F(3,23)=6.6468, p=0.0025] n=6-7. **D)** Freezing levels of animals conditioned with high intensity or not conditioned [Treatment: F (1, 23) = 2.684, p=0.1150. Intensity: F (1, 23) = 57.14, p<0.0001. Interaction: F (1, 23) = 10.58, p=0.0035] n=5-8. *p<0.05 compare to control.

However, since it was observed that TRPV1 channels may be recruited in a intensity-dependent manner (Terzian et al., 2014), we hypothesized that higher levels of intensity may lead to the recruitment of the channel in the dHPC. In this vein, we replicate the experiment using a higher intensity to conditioned the animals (moderate intensity). We observed that 1, 3 or 10nmol of SB administered into the dHPC before retrieval, induced a significant reduction of freezing when compare to vehicle (Fig.1C) [F(3,23)=6.6468, p=0.0025].

In order to further investigate the involvement of intensity in TRPV1 recruitment, we submitted a new subset of mice to higher intensities of conditioning or to an exposure to the CFC apparatus without shock (NC, not-conditioned group). Twenty-four hours later the animals received vehicle or 3nmol of SB administered into the dHPC and were tested for 5min. Our results suggested that, in the absence of conditioning the TRPV1 blocker has no effect on freezing levels, but it impairs retrieval of animals conditioned with the high intense protocol (Fig.1D) [Treatment: F (1, 23) = 2.684, p=0.1150. Intensity: F (1, 23) = 57.14, p<0.0001. Interaction: F (1, 23) = 10.58, p=0.0035]. Taken together these results confirm an intensity-dependent recruitment of dHPC TRPV1 channels in the retrieval of CFC.

### The involvement of anandamide in TRPV1 recruitment in the dHPC

It was previously noticed that after training in the inhibitory avoidance task, AEA levels correlate with the intensity of the training (Morena et al., 2014). Since AEA has 20-times more affinity for CB1 than for TRPV1, more AEA is needed to recruit TRPV1 when CB1 is present in the synapse (Devane et al., 1992; Ross, 2003; Stelt et al., 2005). We hypothesized that the intensity-dependent recruitment of TRPV1 is mediated by the higher levels of AEA released in higher intensities.

In order to probe that SB effect is determine by the balance between AEA-CB1-TRPV1, first we confirmed the co-localization of TRPV1 and CB1 in the dHPC (Fig.2) Second, we quantified the levels of AEA immediately after retrieval in animals submitted to different intensities of conditioning (not-conditioned, low, moderate or high intensity) (Fig.3A-C) [F(3,15)=20.36, p<0,0001]. We observed that there is a correlation between freezing and AEA levels (Fig.3E) [r=0,4707, R=0,2216, p=0.0486] but not 2AG levels (Fig.3E) [r=−0.1618, R=0.02617, p=0.5213]. Next, we pre-treated the animals with a subeffective dose of AM251 (75pmol), a CB1 antagonist, prior to the administration of SB 3nmol (Fig3D). In accordance, the AM251 administration alone was not able to reduce freezing but it prevented the effect of SB (Fig.3E) [Pre-treatment: F(1,28)=0.3097, p=0.5823. Treatment: F (1,28) =4.367, p=0.0458. Interaction: F(1,28) = 1.564, p=0.2214].

**Figure 2.**
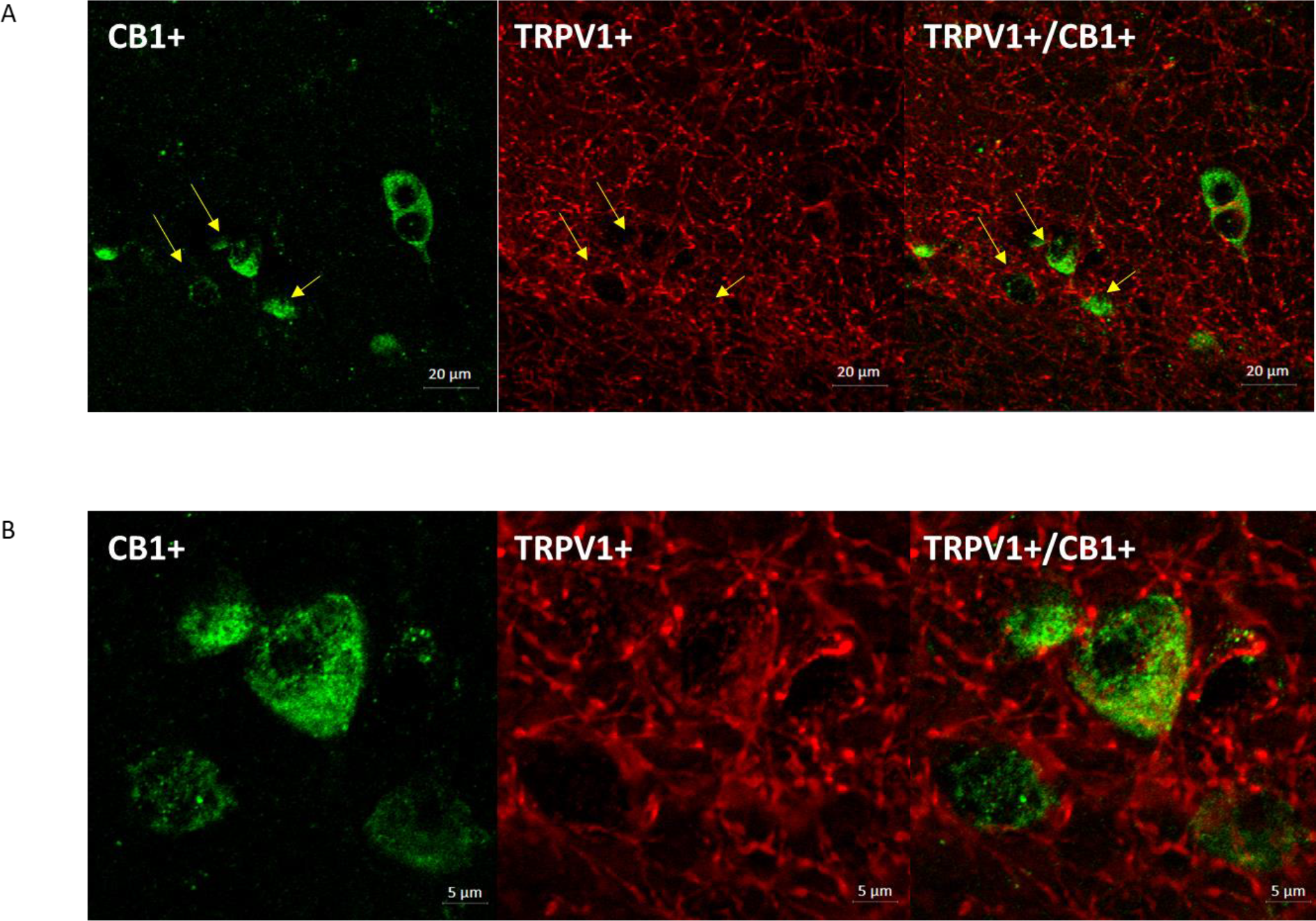
Double immunofluorescence TRPV1-CB1 in the CA1 region of the dHPC. A) Representative image **B)** Augmentation of the representative image showing three TRPV1+ cells colocalizing with CB1. Yellow arrows indicated TRPV1+/CB1+.

**Figure 3.**
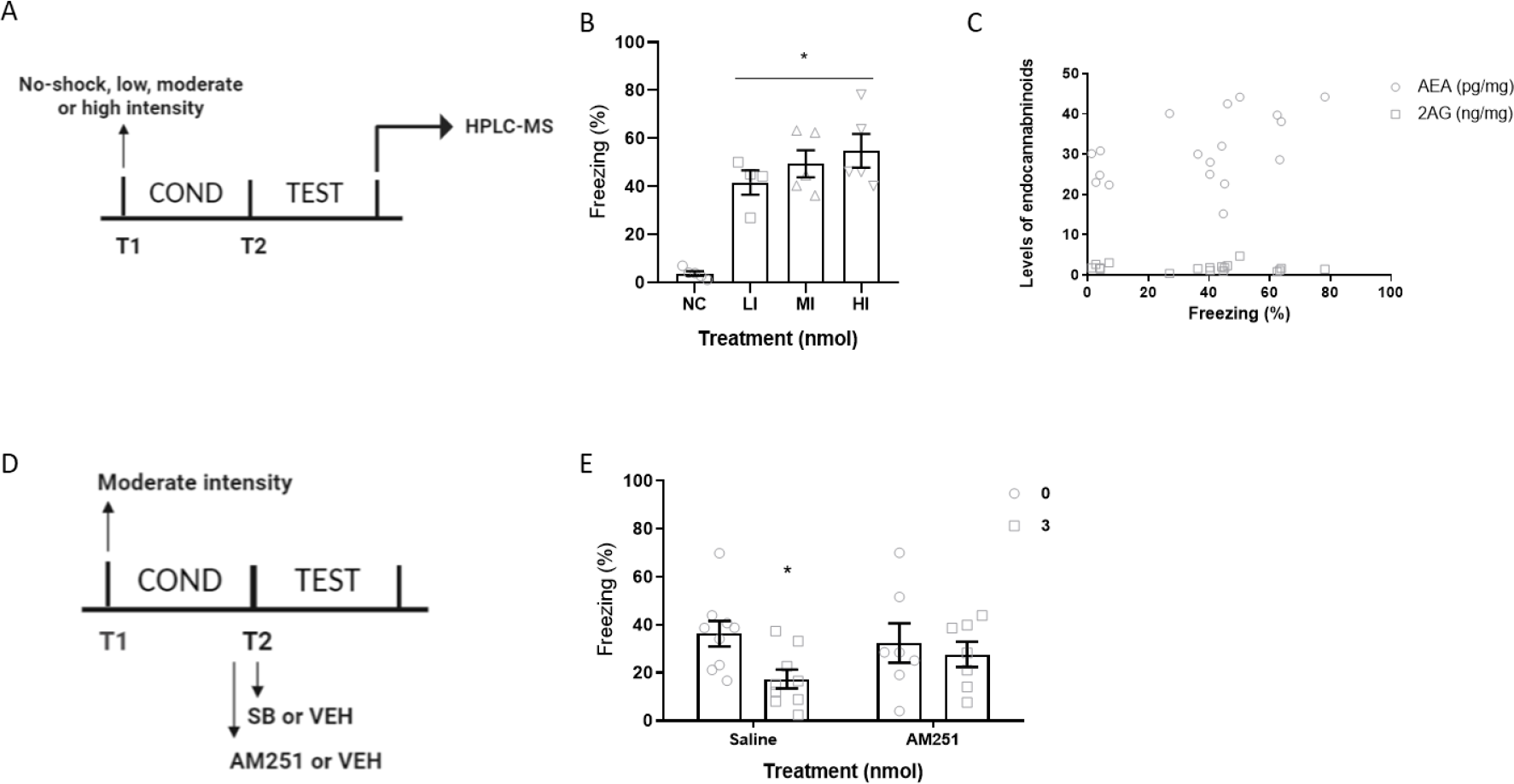
The intensity-dependent recruitment of TRPV1 depends on AEA released. **A)** Experimental design of B and C. **B)** Freezing levels of animals submitted to different intensities of conditioning [F(3,15)=20.36, p<0,0001] n=4-5. **C)** Correlation between freezing levels and endocannabinoid release [AEA, Pearson r=0,4707, R=0,2216, p=0.0486], [2AG, Pearson r=-0.1618, R=0,02617, p=0.5213] n=18. **D)** Experimental design of E. **E)** Pretreatment with subeffective doses of AM251 impairs SB effect [Pre-treatment: F(1,28)=0.3097, p=0.5823. Treatment: F(1,28)=4.367, p=0.0458. Interaction: F(1,28) = 1.564, P=0.2214] n=7-10. *p<0.05 compare to control.

Summarizing, our results suggest that the intensity-dependent recruitment of TRPV1 in the dHPC depends on AEA levels. In addition, SB increases AEA availability to act through CB1 (Fig.4).

**Figure 4.**
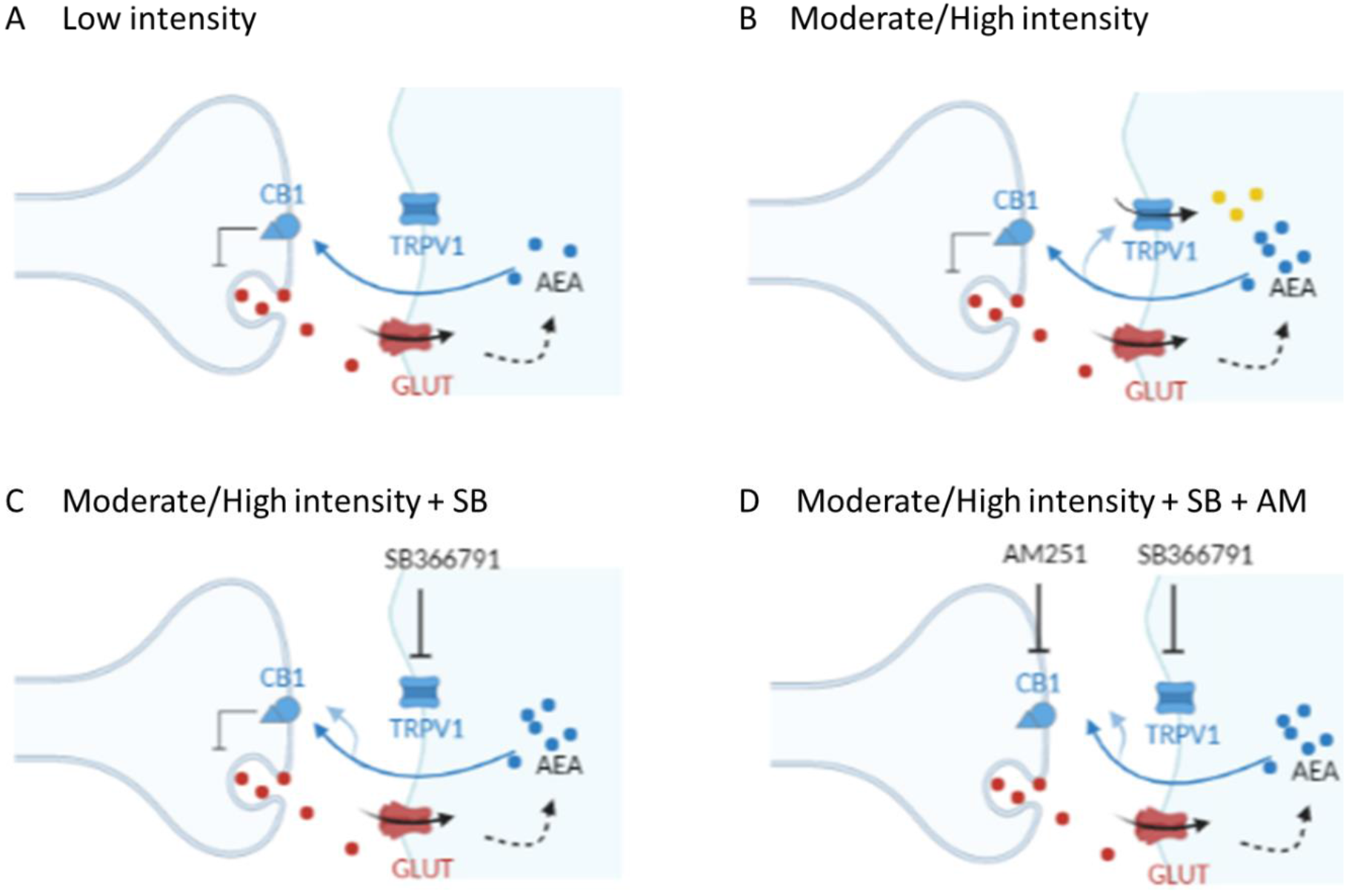
Scheme of TRPV1-AEA-CB_1_ interplay in the retrieval of fear memory. **A)** Retrieval of low-intensity memories: when the conditioning was performed using a low intensity the exposition to the context during retrieval induces a low release of AEA, these levels may be sufficient to recruit CB_1_ receptors but not TRPV1 channels. **B)** Retrieval of moderate and high-intensity memories: when the conditioning was performed using moderate or high intensities, the exposition to the context induced the release of higher levels of AEA, which modulates presynaptic CB_1_ receptors but also postsynaptic TRPV1 channels. C) Retrieval of moderate/high-intensity memories and SB366791 treatment: SB366791 blocks TRPV1 channels increasing AEA availability to act through CB_1_ receptors leading to a decrease in freezing levels. D) Retrieval of moderate/high-intensity memories and a combination of AM251 + SB366791 treatment: a subeffective dose of AM251 prevents the effect of SB366791

### State-dependent effect of dHPC TRPV1 blocking

Several reports suggested that some drugs that interact with fear memory can induced a state-dependent learning, thus, the impairments in freezing responses are not observed when the animal is not under the effects of the drug (Alford & Alford, 1976; Bouton et al., 1990; Colpaert, 1990). In order to address if SB induced an impairment in freezing responses just during their acute effect or it has a sustained effect, we re-test the animals 24h after the treatment with 3nmol of SB. Our results suggested that the treatment effect is sustained 24h later since treated animals did not performed higher levels of freezing in the second test (Fig.5) [Treatment: F (1, 14) = 11.32, p=0.0046. Test: F (1, 14) = 2.320, p=0.1499. Interaction: F (1, 14) = 9.273, p=0.0087].

**Figure 5.**
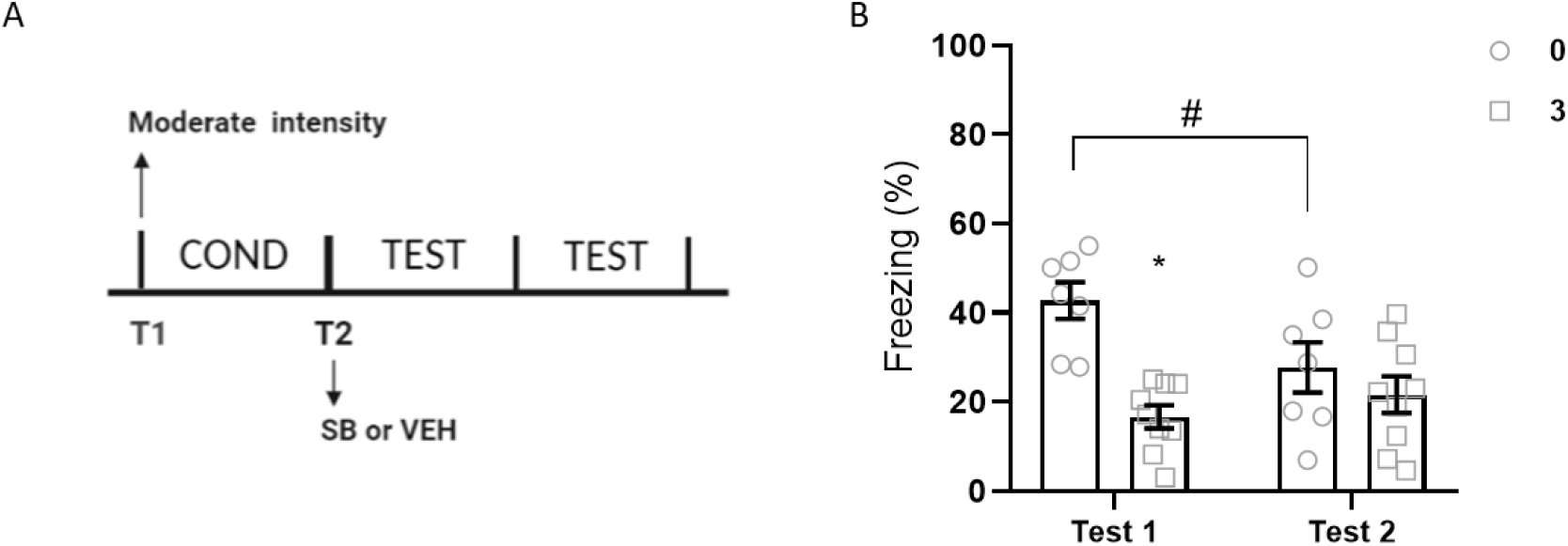
Effect of TRPV1 blocker 24h after administration. **A)** Experimental design. **B)** Animal were treated with SB or vehicle in the retrieval of fear memory (test1) and tested twice, the first 5min after administration and the second 24h later (test2) [Treatment: F (1, 14) = 11.32, p=0.0046. Time: F (1, 14) = 2.320, p=0.1499. Interaction: F (1, 14) = 9.273, p=0.0087] n=7-9. *p<0.05 compare to control. #p<0.05 compare to the same treatment between groups

### Molecular pathways involved in SB366791 effect

Since impairment in retrieval was sustained 24h after the treatment we investigated plasticity pathways potentially involved in this effect. We evaluated the levels of Zif, Arc, TRKB and neurotrophic factors in the HPC of animals conditioned with the moderate intensity and treated with vehicle or 3nmol of SB. The samples were obtained 30min or 24h after the test. In order to assess the effect of surgery and intra-HPC administration, we also evaluated the levels of all the targets 30min after retrieval in animals did not submit to these procedures. Moreover, another group was euthanized and the dHPC dissected 30min after memory acquisition.

#### Plasticity pathways involved in SB366791 effect

First, we assessed the levels of TRPV1, the results suggested that any of the experimental conditions induced changes in TRPV1 RNA. There were not differences between acquisition and retrieval (Fig.S1A) [t=1.178, df=10, p=0,266], neither between animals with or without surgery (Fig.S1B) [t=0.06133, df=9, p=0.9524]. Moreover, we observed similar TRPV1 RNA levels between animals treated with vehicle or SB before retrieval at both time points 30min or 24h after the test (Fig.S1C) [Treatment: F (1,17) = 0.1842, p=0.6732. Time: F (1, 17) = 1,438, p=0.2469. Interaction: F (1,17) = 0.005418, p=0.9422].

Later, we assessed the RNA levels of early genes involved with plasticity, Zif and Arc. Regarding Arc, we did not observed differences between acquisition and retrieval (Fig.6A) [t=0.2572, df=10, p=0,8023] or after retrieval in animals with or without surgery (Fig.S2B) [t=0.6375, df=9, p=0.5397]. However, Zif levels were increased after the acquisition (Fig.6B) [t=3.630, df=9, p=0.0055], but again no differences were observed between animals with or without surgery (Fig.S2C) [t=1.367, df=7, p=0.2139]. In addition, we observed an increase 30 min, but not 24h after retrieval in Arc (Fig.7A) [Treatment: F (1, 19) = 25.9, p<0.0001. Time: F (1, 19) = 33,33, p<0.0001. Interaction: F (1, 19) = 24.77, p<0.0001] and Zif levels (Fig.7B) [Treatment: F (1, 16) = 87,48, p<0.0001. Time: F (1, 16) = 147,5, p<0.0001. Interaction: F (1, 16) = 89,43, p<0.0001] in animals treated with 3nmol of SB.

**Figure 6.**
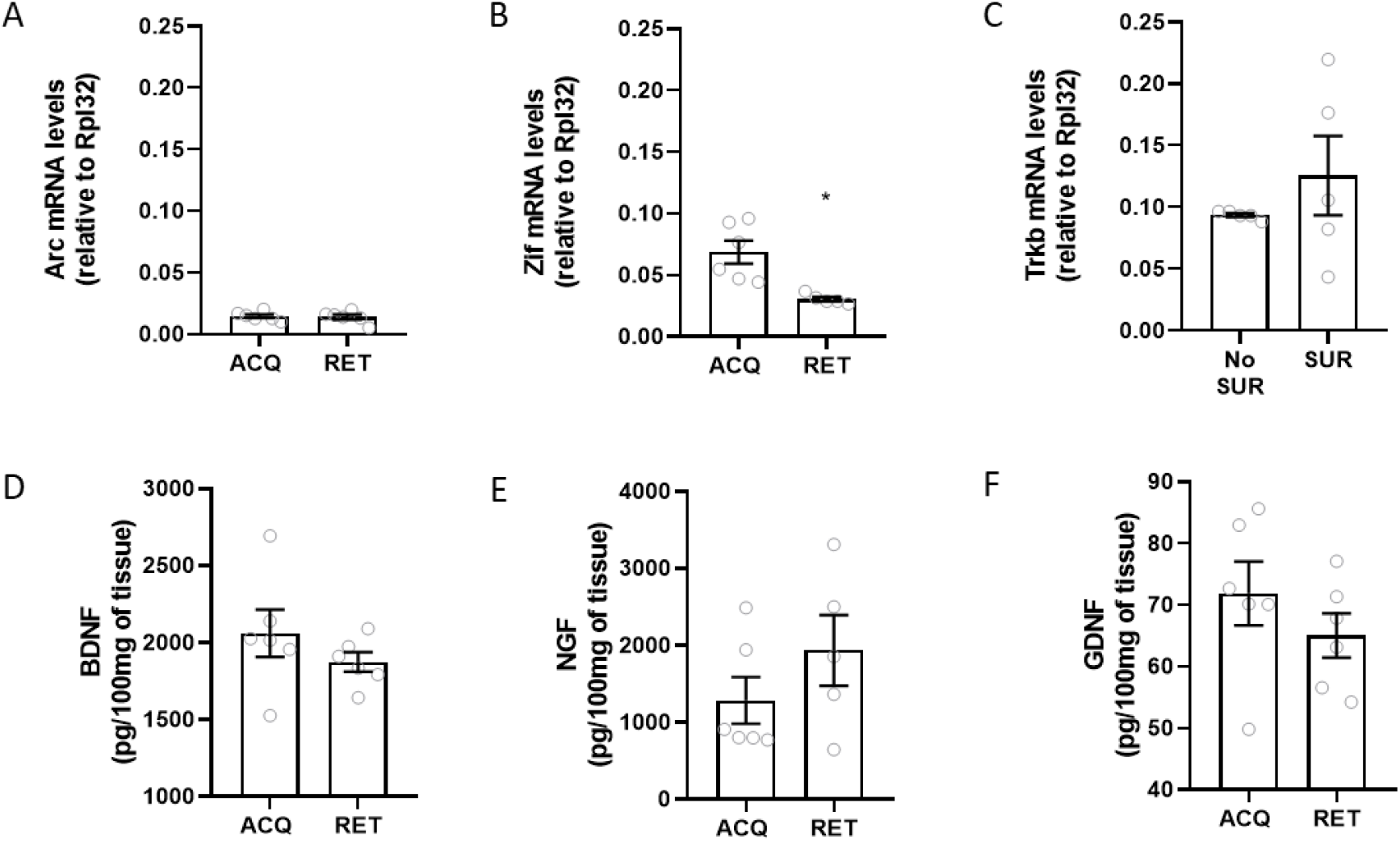
Early genes and neurotrophic factors 30min after acquisition or retrieval in animals conditioned with moderate intensity. **A)** Arc mRNA levels [t=0.2572, df=10, p=0,8023] n=6. **B)** Zif mRNA levels [t=3.630, df=9, p=0.0055] n=5-6. **C)** Trkb mRNA levels [t=2.055, df=9, p=0,0700] n=5-6. **D)** BDNF levels [t=1.116, df=10, p=0,2904] n=6. **E)** NGF levels [t=1.221, df=9, p=0.2530] n=5-6. **F)** GDNF levels [t=1.087, df=10, p=0.3027] n=6. *p<0.05 compare to control.

**Figure 7.**
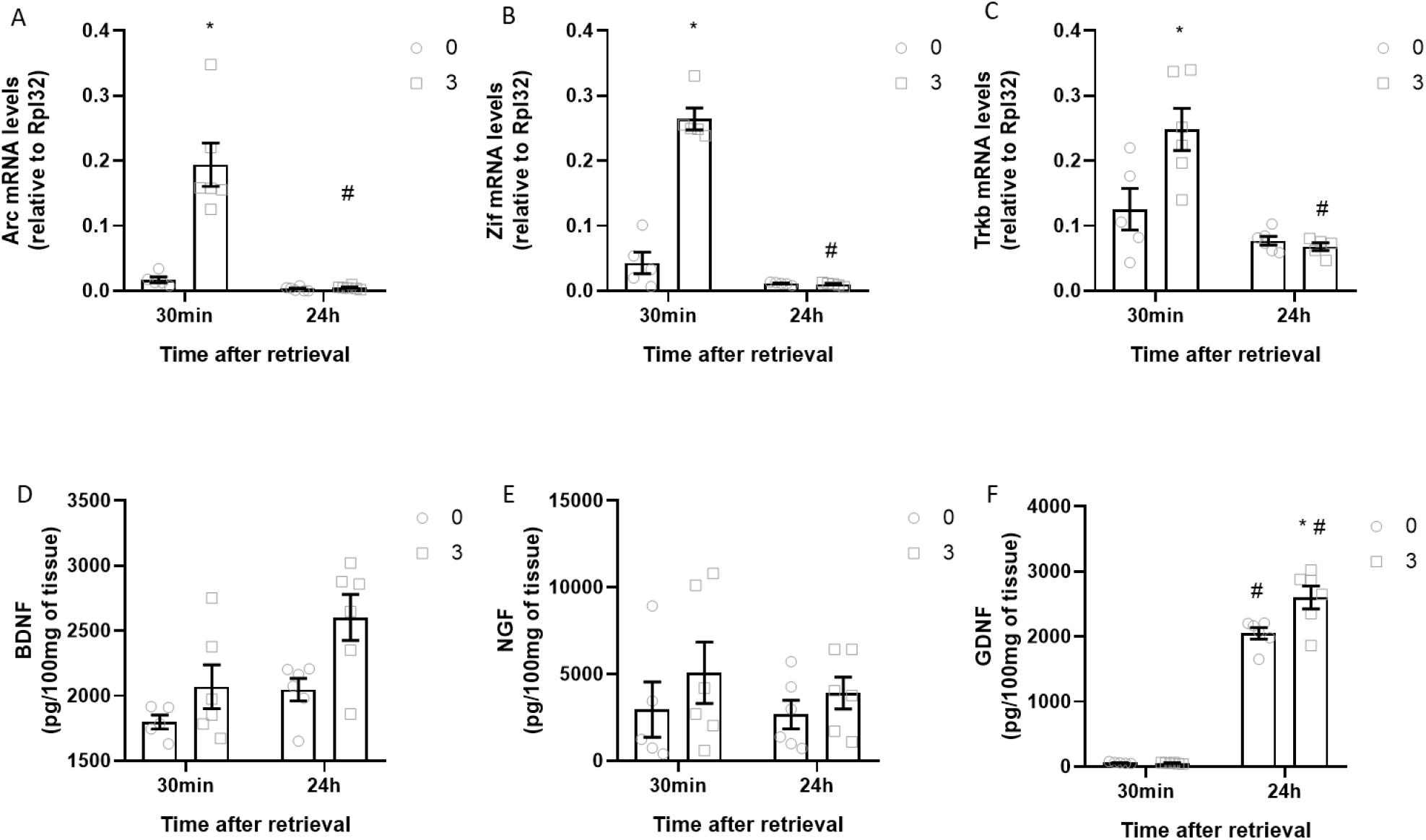
Early genes and neurotrophic factors 30min or 24h after retrieval in animals conditioned with the moderate intensity**. A)** Arc mRNA levels [Treatment: F (1, 19) = 25.9, p<0.0001. Time: F (1, 19) = 33,33, p<0.0001. Interaction: F (1, 19) = 24.77, p<0.0001] n=5-6**. B)** Zif mRNA levels [Treatment: F (1, 16) = 87,48, p<0.0001. Time: F (1, 16) = 147,5, p<0.0001. Interaction: F (1, 16) = 89,43, p<0.0001] n=5-6**. C)** Trkb mRNA levels [Treatment: F (1,18) = 5.836, p=0.0265. Time: F (1, 18) = 23.81, p=0.0001. Interaction: F (1, 18) =7.946, p=0.0114] n=5-6**. D)** BDNF levels [Treatment: F (1,19) = 8.946 p=0.0075. Time: F (1, 19) = 8.035, p=0.0106, Interaction: F (1,19) = 1.059, p= 0.3164] n=5-6. **E)** NGF levels [Treatment: F (1,19) = 1.610 p=0.2198. Time: F (1, 19) = 0.2969, p=0.5922, Interaction: F (1,19) = 0.1121, p= 0.7414] n=5-6**. F)** GDNF levels [Treatment: F (1,19) = 7.061 p=0.0156. Time: F (1, 19) = 483.1, p<0.0001, Interaction: F (1,19) = 7.408, p= 0.0135] n=5-6. *p<0.05 compare to control. #p<0.05 compare to the same treatment between groups.

Finally, we investigated the involvement of neurotrophic factors. First, we assessed the RNA levels of Trkb receptors, and we observed a similar pattern compared to Arc: no differences were identified between acquisition and retrieval (Fig.6C) [t=2.055, df=9, p=0,0700], or after the retrieval in animals with and without surgery (Fig.S2D) [t=0.9955, df=8, p=0.3486]; however, the animals treated with 3nmol of SB presented and increase in Trkb RNA 30min but not 24h after the test (Fig7A) [Treatment: F (1,18) = 5.836, p=0.0265. Time: F (1, 18) = 23.81, p=0.0001. Interaction: F (1, 18) =7.946, p=0.0114].

In order to assessed the levels of neurotrophic factors, the samples were analysed by ELISA. We did not find differences between acquisition and retrieval in BDNF (Fig.6D) [t=1.116, df=10, p=0,2904], NGF (Fig.6E) [t=1.221, df=9, p=0.2530] or GDNF (Fig.6F) [t=1.087, df=10, p=0.3027]. Similarly, there were no differences between animals with or without surgery in any of the neurotrophic factors: BDNF (Fig.S2E) [t=0.816, df=9, p=0.3958], NGF (Fig.S2F) [t=0.6176, df=8, p=0.5540] or GDNF (Fig.S2G) [t=0.7230, df=9, p=0.4881].

Analysis at different time points of BDNF expression indicated that animals treated with SB 3nmol presented higher levels of this neurotrophic factor 30min after retrieval, and those levels were higher after 24h (Fig.7D) [Treatment: F (1,19) = 8.946 p=0.0075. Time: F (1, 19) = 8.035, p=0.0106, Interaction: F (1,19) = 1.059, p= 0.3164]. Regarding NGF, we did not observed differences among treatments and time points (Fig.7D) [Treatment: F (1,19) = 1.610 p=0.2198. Time: F (1, 19) = 0.2969, p=0.5922, Interaction: F (1,19) = 0.1121, p= 0.7414]. Finally, GDNF levels seemed increased 24h after the test, and those levels were higher in the group treated with SB (Fig.7E) [Treatment: F (1,19) = 7.061 p=0.0156. Time: F (1, 19) = 483,1, p<0.0001, Interaction: F (1,19) = 7.408, p= 0.0135].

#### Neuroimmune markers involved in SB366791 effect

Recently reports indicated that CFC is able to orchestrate changes in microglia morphology (Chaaya et al., 2019), suggesting the involvement of these cells in emotional memory (Enomoto & Kato, 2021). Since, TRPV1 channels may be expressed in microglia (Marrone et al., 2017) and we observed changes in the levels of GDNF we asked ourselves if TRPV1 blockers may recruit other components of the immune network in the dHPC.

First, we evaluated the co-localization of TRPV1 channels and NeuN, our results suggested that not all the cells TRPV1^+^ in the dHPC are neurons (Fig.8). Later, we evaluated by PCR the RNA levels of Cx3Cr1. We did not observe any differences between acquisition and retrieval (Fig.9A) [t=1.028, df=10, p=0,3280] or after retrieval in animals with or without surgery (Fig. S2H) [t=0.1373, df=9, p=0.8938]. Similarly, to the pattern observed in plasticity factors, Cx3Cr1 RNA was increased in the treated group 30min but not 24h after the test (Fig.10A) [Treatment: F (1,18) = 4.141, p=0.0569. Time: F (1, 18) = 1.696, p=0.2092. Interaction: F (1,18) = 6.566, p=0.0196]. To further investigate the involvement of neuroimmune responses, we evaluated by ELISA the levels of CX3CL1, IL-1β, IL-10, IL-6 and TNFα. Despite the surgery was performed 8 days before the test and in accordance with the invasive character of the surgery procedure, animals submitted to it presented higher levels of all the targets studied when compare to animals without surgery: CXCL1 (Fig. S2I) [t=4.507, df=9, p=0.0015], IL-1β (Fig. S2J) [t=6.117, df=9, p=0.0002], IL-6 (Fig. S2K) [t=3.959 df=9, p=0.0033], IL-10 (Fig. S2L) [t=3.727, df=9, p=0.0047], and TNFα (Fig. S2M) [t=3.052 df=9, p=0.0137].

**Figure 8.**
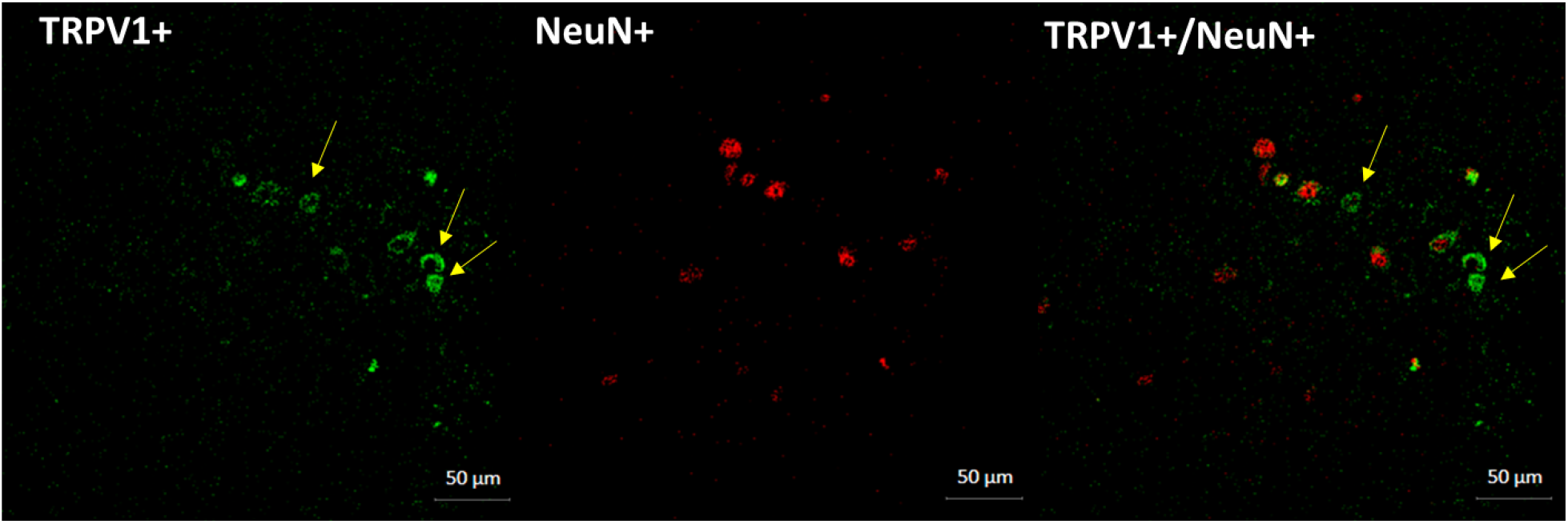
Double immunofluorescence TRPV1-NeuN in the CA1 region of the dHPC. Yellow arrows indicate TRPV1+ not co-localized with NeuN+

**Figure 9.**
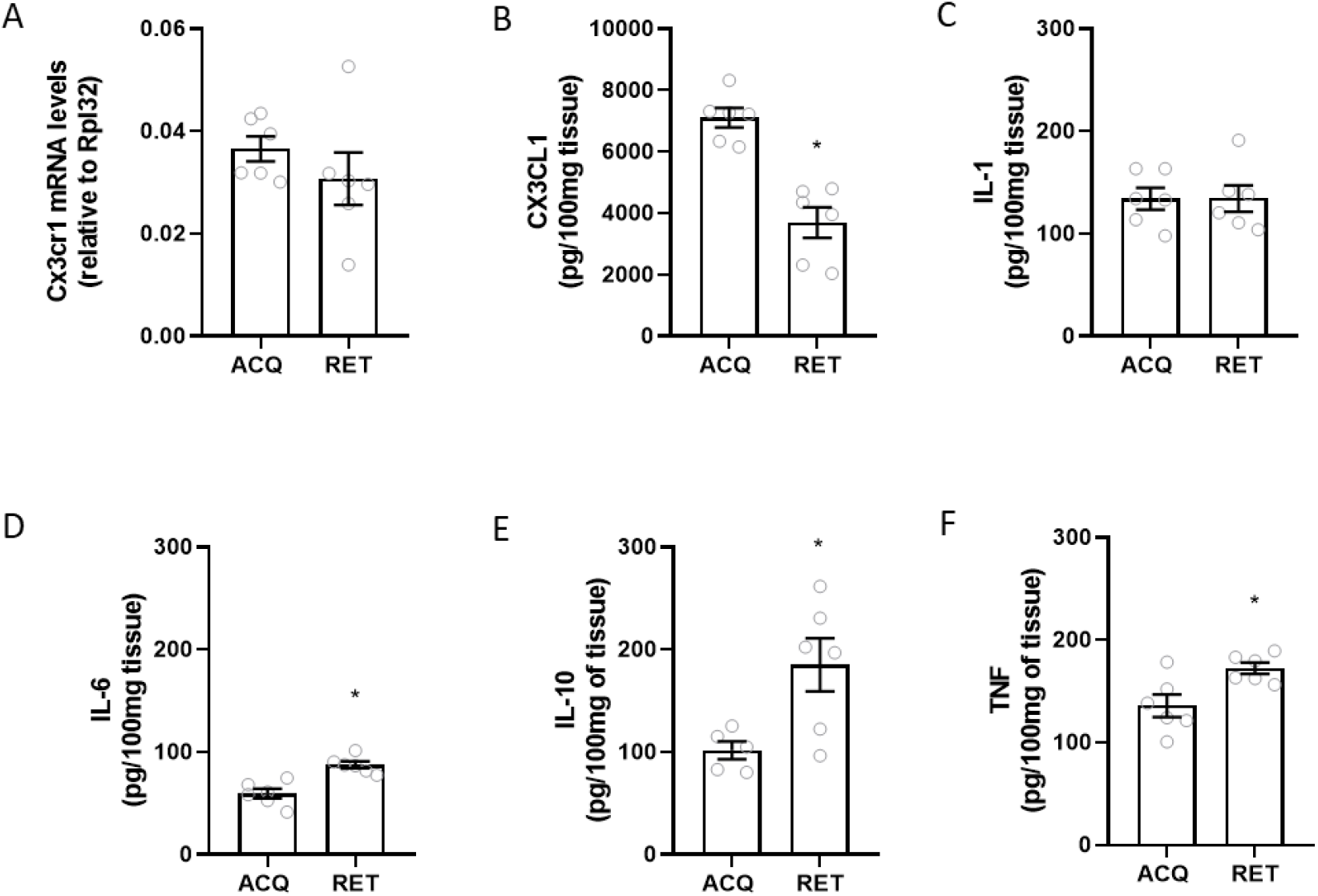
Neuroinflammatory response 30min after acquisition or retrieval in animals conditioned with moderate intensity. **A)** Cx3cr1 mRNA levels [t=1.028, df=10, p=0.3280] n=6. **B)** CX3CL1 levels [t=5.791, df=10, p=0.0002] n=6. **C)** IL-1β levels [t=0.006285, df=10, p=0.9951] n=6. **D)** IL-6 levels [t=4.808, df=10, p=0.0007] n=6. **E)** IL-10 levels [t=2.803, df=9, p=0.0206] n=5-6. **F)** TNFα levels [[t=2.952, df=10, p=0.0145] n=6. *p<0.05 compare to control.

**Figure 10.**
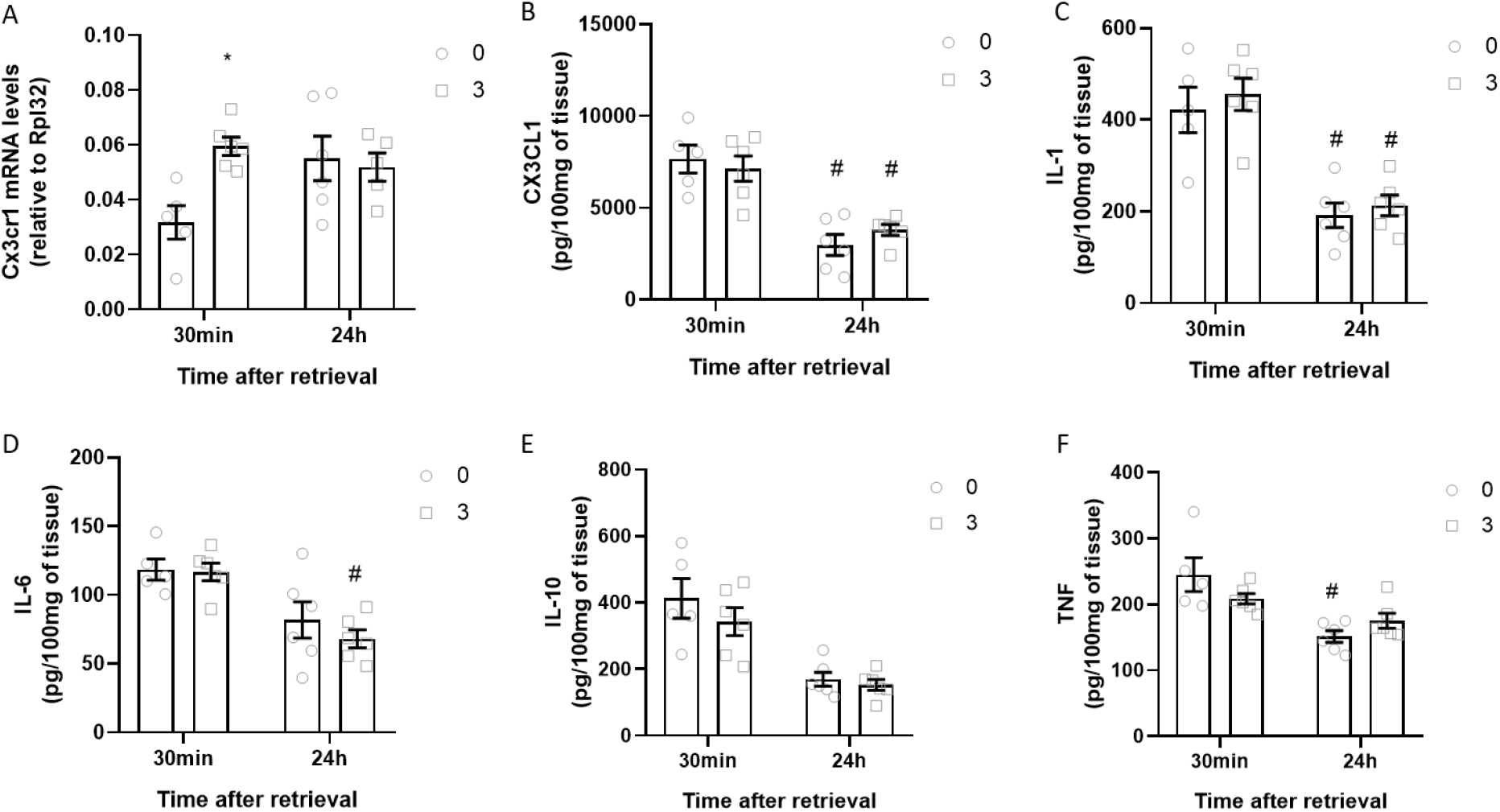
Neuroinflammatory response 30min or 24h after retrieval in animals conditioned with the moderate intensity. **A)** Cx3cr1 mRNA levels [Treatment: F (1,18) = 4.141, p=0.0569. Time: F (1, 18) = 1.696, p=0.2092. Interaction: F (1,18) = 6.566, p=0.0196] n=5-6. **B)** CX3CL1 levels [Treatment: F (1,19) = 0.06, p=0.8091. Time: F (1, 19) = 45.54, p<0.0001. Interaction: F (1,19) = 1.299, p=0.2685] n=5-6. **C)** IL-1β levels [Treatment: F (1,19) = 0.6733, p=0.422. Time: F (1, 19) = 49.06, p<0.0001. Interaction: F (1,19) = 0.03912, p=0.8453] n=5-6. **D)** IL-6 levels [Treatment: F (1,19) = 0.7345 p=0.4021. Time: F (1, 19) = 22.23, p=0.0002. Interaction: F (1,19) = 0.4346, p=0.5177] n=5-6**. E)** IL-10 levels [Treatment: F (1,19) = 1.426 p=0.2471. Time: F (1, 19) = 35.12, p<0.0001. Interaction: F (1,19) = 0.5392, p=0.4717] n=5-6. **F)** TNFα levels [Treatment: F (1,19) = 0.2093 p=0.6525. Time: F (1, 19) = 20.92, p=0,0002. Interaction: F (1,19) = 4.682, p= 0.0434] n=5-6. *p<0.05 compare to control. #p<0.05 compare to the same treatment between groups.

Moreover, no differences were observed between retrieval and acquisition regarding IL-1β levels (Fig. 9B) [t=0.006285, df=10, p=0.9951]. However, lower levels of IL6 levels (Fig. 9D) [t=4.808, df=10, p=0.0007], IL-10 (Fig. 9E) [t=2.803, df=9, p=0.0206], and TNFα (Fig. 9F) [t=2.952, df=10, p=0.0145] were founded in the acquisition group compare to the retrieval group. In contrast, higher levels of CX3CL1 (Fig. 9B) [t=5.791, df=10, p=0.0002] were observed in the acquisition group, suggesting different involvement of neuro-inflammatory markers among phases of CFC.

Finally, we observed a decreased in all the targets evaluated 24h after the test independent of treatment: CX3CL1 (Fig. 10B) [Treatment: F (1,19) = 0.06, p=0.8091. Time: F (1, 19) = 45.54, p<0.0001. Interaction: F (1,19) = 1.299, p=0.2685], IL-1β (Fig. 10C) [Treatment: F (1,19) = 0.6733, p=0.422. Time: F (1, 19) = 49.06, p<0.0001. Interaction: F (1,19) = 0.03912, p=0.8453], IL-6 (Fig. 10D) [Treatment: F (1,19) = 0.7345 p=0.4021. Time: F (1, 19) = 22.23, p=0.0002. Interaction: F (1,19) = 0.4346, p=0.5177], IL-10 (Fig. 10E) [Treatment: F (1,19) = 1.426 p=0.2471. Time: F (1, 19) = 35.12, p<0.0001. Interaction: F (1,19) = 0.5392, p=0.4717] and TNFα (Fig. 10F) [Treatment: F (1,19) = 0.2093 p=0.6525. Time: F (1, 19) = 20.92, p=0,0002. Interaction: F (1,19) = 4.682, p= 0.0434].

### Molecular pathways involved in SB366791 effect are biased by intensity

We observed that TRPV1 channels recruitment depends on conditioning intensity, in this sense, our results showed that SB impairs the retrieval of fear memory when the animals are conditioned with moderate or high intensity but not with low intensity. Since intensity seems to be a key element in TRPV1 recruitment, we explored the involvement of this factor in biased the molecular pathways recruited by SB in intensities where this blocker was able to impair retrieval; moderate intensity (MI) and high intensity (HI). Since the dHPC is involved in processing and storage spatial memory (Eichenbaum et al., 1999; Moser et al., 2015), we include a control group of not-conditioned memory, this group receive the treatment and was exposed to the CFC apparatus but it was not submitted to the shock (NC). The three groups were treated with vehicle or SB 5min before the test and the HPC was dissect 30min after.

#### Intensity as a regulator of plasticity pathways involved in SB366791 effect

First, we compared the mRNA levels of TRPV1 in the HPC, we observed no effect of intensity or treatment (Fig. S1D) [Treatment: F (1, 26) = 2.978, p=0.0963. Intensity: F (2,26) = 1.254, p=0.3021. Interaction: F (2, 26) = 0.6268, p=0.5422]. In contrast, Arc RNA is increased due to treatment only in animals conditioned with moderate intensity, but not in animals not submitted to shock or conditioned with the high intensity protocol (Fig. 11A) [Treatment: F (1, 27) = 23.15, p<0.0001. Intensity: F (2,27) = 8,182, p=0.0017. Interaction: F (2, 27) = 11.50, p=0,0002]. On the other hand, Zif RNA was increased by the treatment in animals not-conditioned or conditioned with the moderate intensity, but again this was not observed in animals conditioned with high intensities (Fig. 11B) [Treatment: F (1, 25) = 60.53, p<0.0001. Intensity: F (2,25) = 11.30, p=0.0003. Interaction: F (2, 25) = 11.34, p=0.0003]. The pattern behind Trkb RNA levels was similar to that observed in Arc: the results suggested that the increase induced by the treatment is associated with moderate intensities of conditioning, since no differences between vehicle and treated groups were observed in animals not-conditioned or conditioned with high intensity (Fig. 11C) [Treatment: F (1, 27) = 5,550, p=0.0260. Intensity: F (2,27) = 4.311, p=0.0237. Interaction: F (2, 27) = 3.480, p=0.0452]. In the same vein, levels of BDNF were significantly affected by the intensity of the protocol (Fig. 11D) [Treatment: F (1,27) = 1.877, p=0.1820. Intensity: F (2,27) = 6.823, p=0.0040, Interaction: F (2, 27) = 0.7236, p=0.4942]. However, neither intensity nor treatment induced significant changes on NGF (Fig. 11E) [Treatment: F (1,27) = 1. 520, p=0.2283. Intensity: F (2,27) = 0.07576, p=0.9242, Interaction: F (2, 27) =0.3695, p=0.6945] or GDNF levels (Fig. 11F) [Treatment: F (1,27) = 0.6003, p=0.4452. Intensity: F (2,27) = 1.830, p=0.1797, Interaction: F (2, 27) =0.4419 p=0.6474].

**Figure 11.**
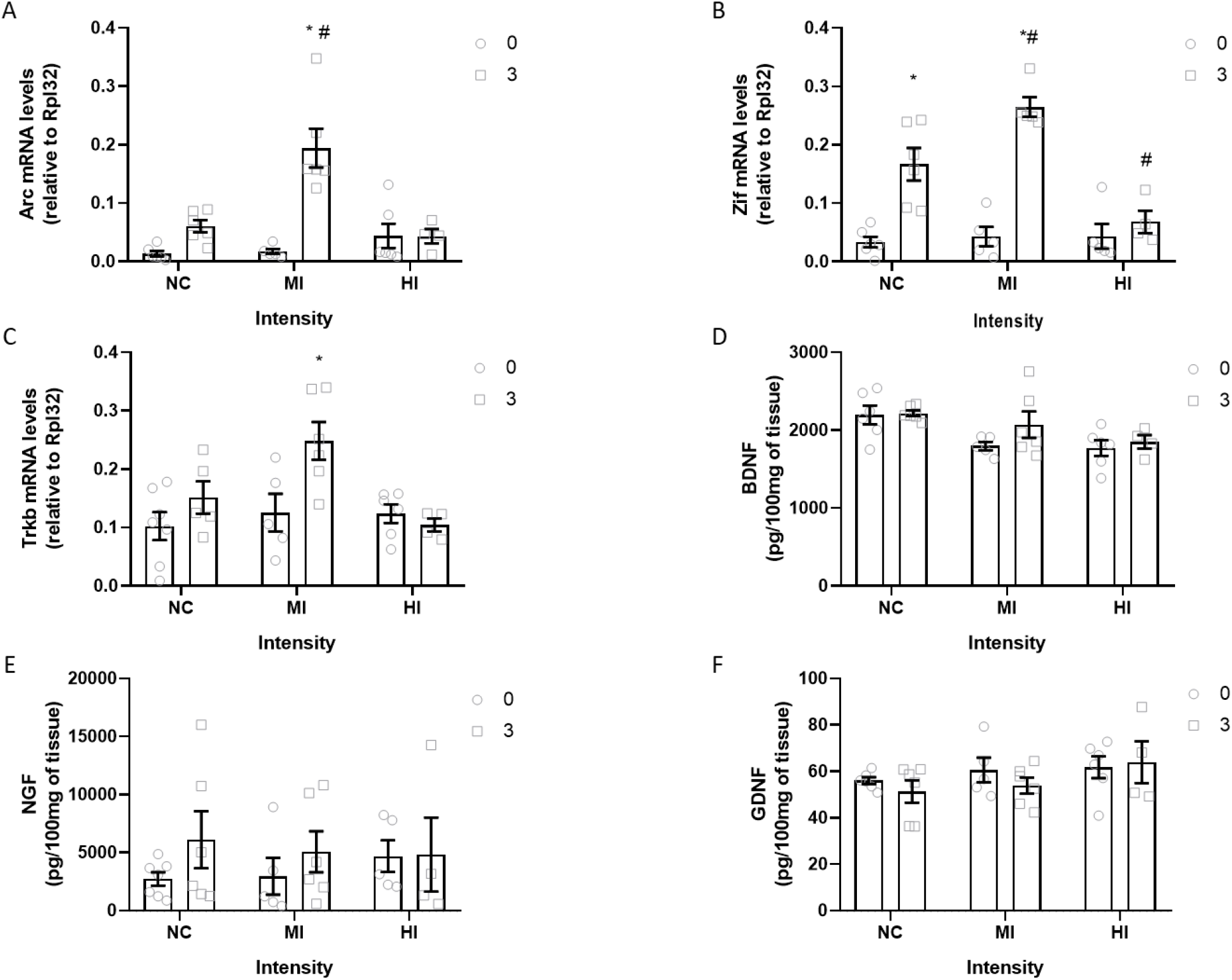
Early genes and neurotrophic factors 30min after retrieval in animals conditioned with the moderate intensity (MI), high intensity (HI) or not-conditioned (NC). **A)** Arc mRNA levels [Treatment: F (1, 27) = 23.15, p<0.0001. Intensity: F (2,27) = 8,182, p=0.0017. Interaction: F (2, 27) = 11.50, p=0.0002] n=5-6. **B)** Zif mRNA levels [Treatment: F (1, 25) = 60.53, p<0.0001. Intensity: F (2,25) = 11.30, p=0.0003. Interaction: F (2, 25) = 11.34, p=0.0003] n=4-6. **C)** Trkb mRNA levels [Treatment: F (1, 27) = 5,550, p=0.0260. Intensity: F (2,27) = 4.311, p=0.0237. Interaction: F (2, 27) = 3.480, p=0.0452] n=4-7. **D)** BDNF [Treatment: F (1,27) = 1.877, p=0.1820. Intensity: F (2,27) = 6.823, p=0.0040, Interaction: F (2, 27) = 0.7236, p=0.4942] n=4-7. **E)** NGF levels [Treatment: F (1,27) = 1. 520, p=0.2283. Intensity: F (2,27) = 0.07576, p=0.9242, Interaction: F (2, 27) =0.3695, p=0.6945] n=4-7. **F)** GDNF levels [Treatment: F (1,27) = 0.6003, p=0.4452. Intensity: F (2,27) = 1.830, p=0.1797, Interaction: F (2, 27) =0.4419 p=0.6474] n=4-6. *p<0.05 compare to control. #p<0.05 compare to the same treatment between groups.

#### Intensity as a regulator of neuroimmune markers involved in SB366791 effect

Regarding Cx3cr1, we found no differences among intensities (Fig. 12A) [Treatment: F (1, 28) = 13.65, p=0.0009. Intensity: F (2,28) = 0.6041, p=0.5536. Interaction: F (2, 28) = 2.067, p=0.1455]. Curiously, despite all the animals were submitted to surgery and tested 8 days before, the results suggested that the expression levels of all the targets addressed is modulated by the intensity of the conditioning but not by the treatment: CX3CL1 (Fig. 12B) [Treatment: F (1, 27) = 0.2970, p=0.5902. Intensity: F (2,27) = 6.832, p=0.004. Interaction: F (2, 27) = 0.2412, p=0.7873], IL-1β (Fig. 12C) [Treatment: F (1, 27) = 3.010, p=0.0941. Intensity: F (2,27) = 10.98, p=0.0003. Interaction: F (2, 27) = 0.3165, p=0.7313], IL-6 (Fig. 12D) [Treatment: F (1,26) = 2.253 p= 0.1454. Intensity: F (2,26) = 15.64, p<0,0001. Interaction: F (2, 26) = 2.412, p= 0.1094], IL-10 (Fig. 12E) [Treatment: F (1,28) = 1.363, p=0.2529. Intensity: F (2,28) = 17.80, p<0,0001. Interaction: F (2, 28) = 3.188, p=0.0.0566], and TNFα (Fig. 12F) [Treatment: F (1,27) = 0.04761, p=0.8289. Intensity: F (2,27) = 6.949, p=0.0037. Interaction: F (2, 27) = 3.3254, p=0.0542]. These results suggest that the immune response induced by the surgery is modulated by the intensity of conditioning.

**Figure 12.**
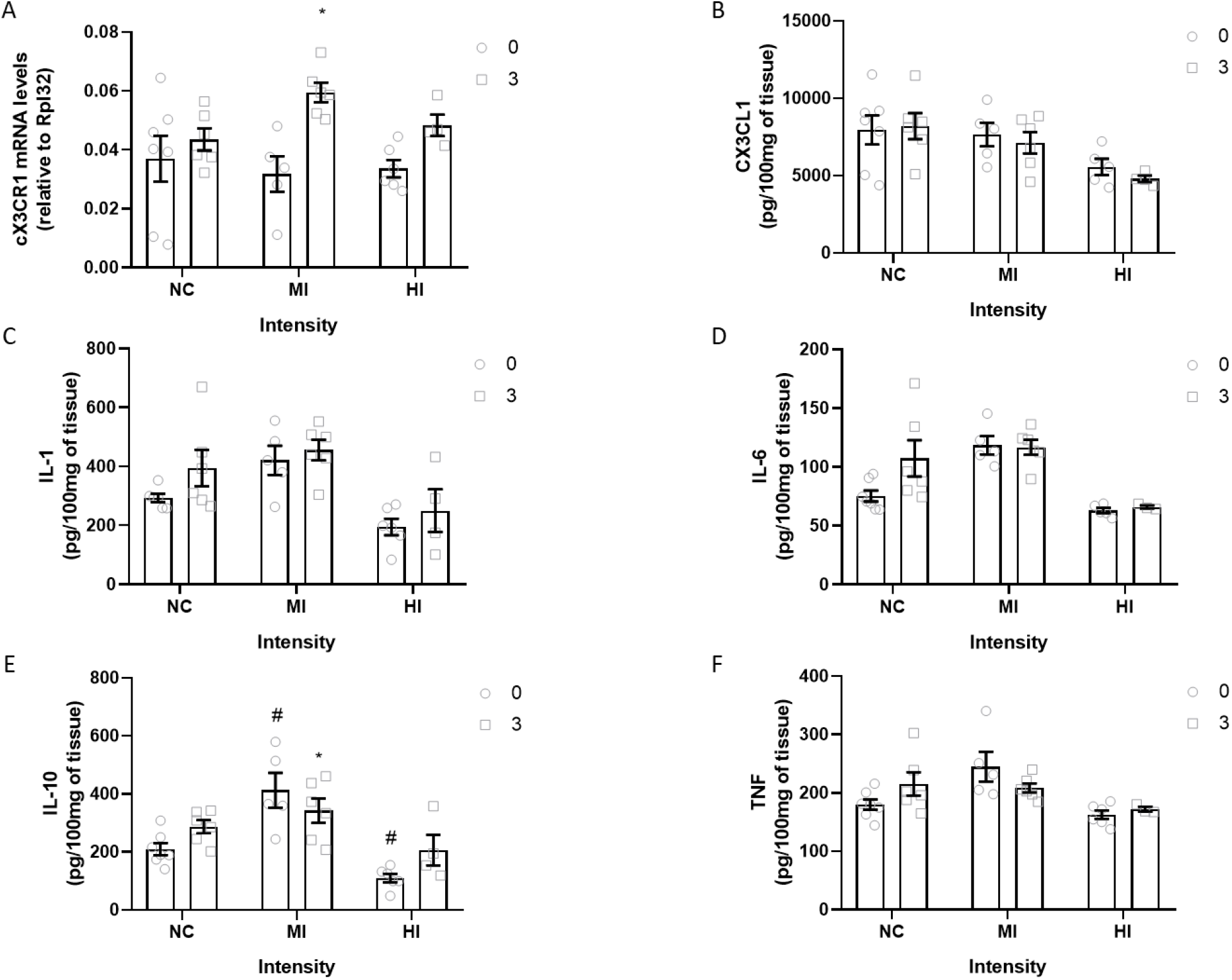
Neuroinflammatory response 30min after retrieval in animals conditioned with the moderate intensity (MI), high intensity (HI) or not-conditioned (NC). **A)** Cx3cr1 mRNA levels [Treatment: F (1, 28) = 13.65, p=0.0009. Intensity: F (2,28) = 0.6041, p=0.5536. Interaction: F (2, 28) = 2.067, p=0.1455] n=5-6. **B)** CX3CL1 levels [Treatment: F (1, 27) = 0.2970, p=0.5902. Intensity: F (2,27) = 6.832, p=0.004. Interaction: F (2, 27) = 0.2412, p=0.7873] n=4-7. **C)** IL-1β levels [Treatment: F (1, 27) = 3.010, p=0.0941. Intensity: F (2,27) = 10.98, p=0.0003. Interaction: F (2, 27) = 0.3165, p=0.7313] n=4-6. **D)** IL-6 levels [Treatment: F (1,26) = 2.253 p= 0.1454. Intensity: F (2,26) = 15.64, p<0,0001. Interaction: F (2, 26) = 2.412, p= 0.1094] n=3-7. **E)** IL-10 levels [Treatment: F (1,28) = 1.363, p=0.2529. Intensity: F (2,28) = 17.80, p<0,0001. Interaction: F (2, 28) = 3.188, p=0.0566] n=4-6. **F)** TNFα levels [Treatment: F (1,27) = 0.04761, p=0.8289. Intensity: F (2,27) = 6.949, p=0.0037. Interaction: F (2, 27) = 3.3254, p=0.0542] n=3-5. *p<0.05 compare to control. #p<0.05 compare to the same treatment between groups.

### The involvement of intensity in long-term effects of SB

Our results indicated that different intensities of conditioning may biased the recruitment of molecular pathways by SB without preventing the impairment in memory retrieval. In order to evaluated if this bias in molecular pathways may have any effect in memory after retrieval, we evaluated the extinction and reinstatement in animals conditioned with moderate and high intensity. First the animals were conditioned with one of the intensities, 24h later they were treated with the blocker or vehicle and tested for 5min. Later, they were submitted to an extinction and reinstatement sessions and finally re-tested (Fig. 13A).

**Figure 13.**
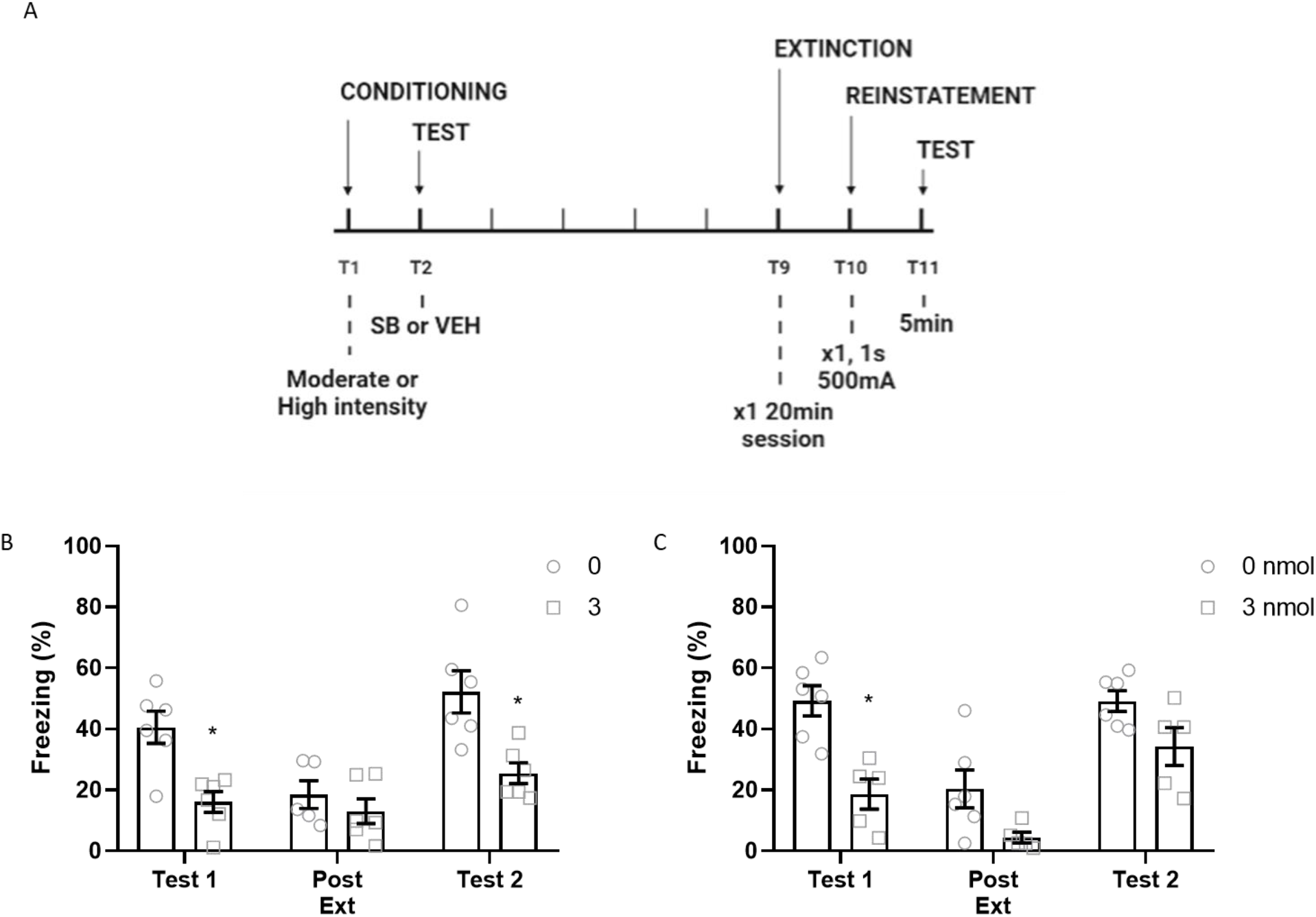
Freezing levels of animals treated with vehicle or SB 3nmol before retrieval during the test 1, after extinction and test 2. **A)** Experimental design**. B)** Animals conditioned with moderate intensity [Treatment: F (1,29) = 23,40, p>0,0001. Time: F (2,29) = 11,41, p=0.0002. Interaction: F (2,29) = 2.888, p=0.0718] n=5-6**. C)** Animals conditioned with high intensity [Treatment: F (1,27) = 25,94, p<0,0001. Time: F (2,27) = 18.96, p<0.0001. Interaction: F (2.27) = 1,574, p=0,2256] n=5-6. *p<0.05 compare to control.

**Figure 14:**
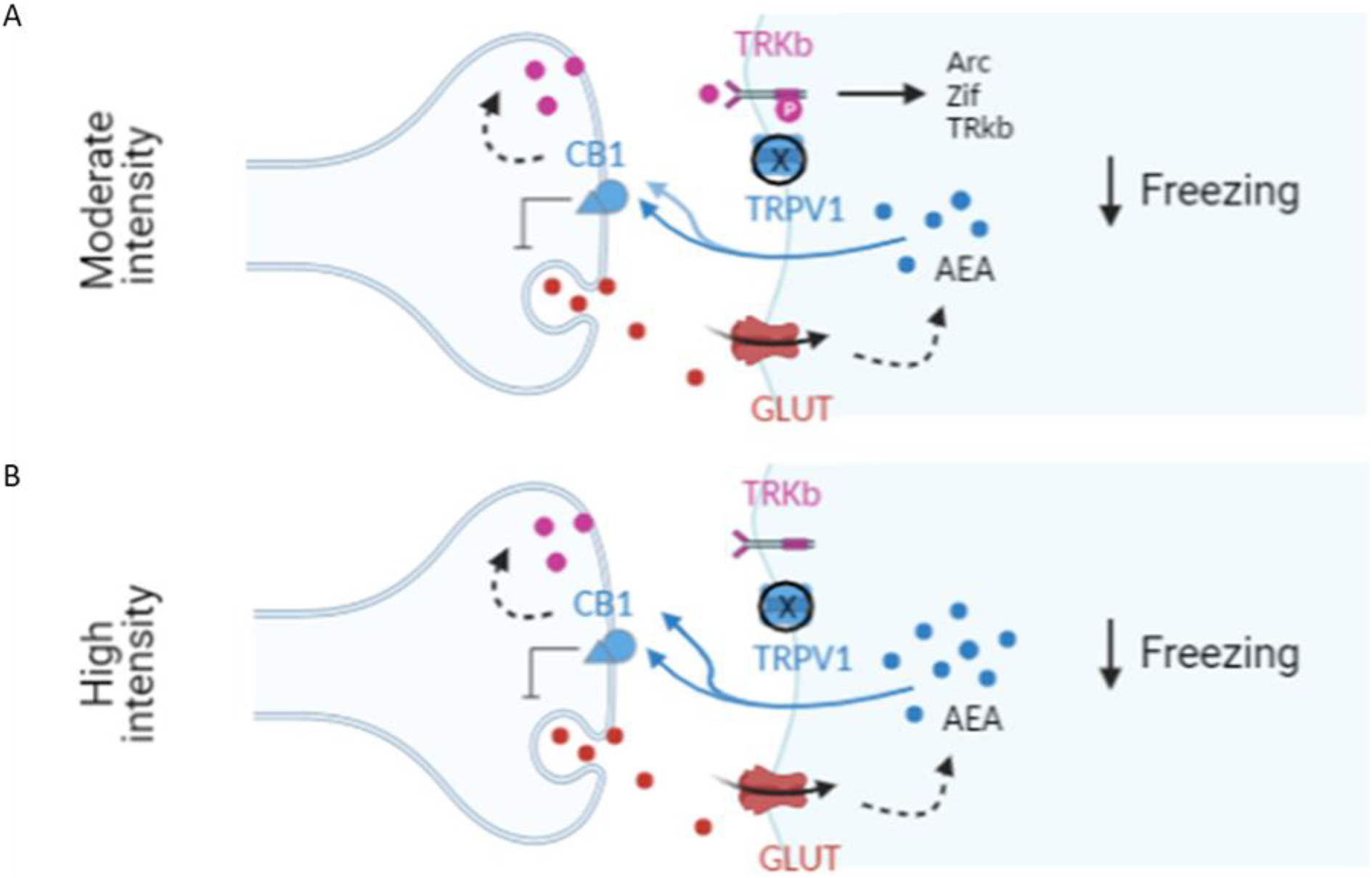
Intensity biased the molecular pathways recruited after blocking TRPV1.

As expected, in the first test the TRPV1 blocker was able to decrease freezing in animals conditioned with the moderate (Fig. 13B) and the high intensity (Fig. 13C). Moreover, we observed that the levels of freezing in the control group decreased after extinction achieving similar levels to those observed in the treated group in both, moderate and high intensity. However, in the second test, animals conditioned with the moderate intensity and treated with the TRPV1 blocker in the first test were not affected by the reinstatement session (Fig. 12B) [Treatment: F (1,29) = 23,40, p>0,0001. Time: F (2,29) = 11,41, p=0.0002. Interaction: F (2,29) = 2.888, p=0.0718], but no differences were observed between vehicle and treatment groups in animals conditioned with the high intensity protocol (Fig. 12C) [Treatment: F (1,27) = 25,94, p<0,0001. Time: F (2,27) = 18.96, p<0.0001. Interaction: F (2.27) = 1,574, p=0,2256]. The freezing levels observed during the second test cannot be explain by the intensity of the shock used during the reinstatement session (Fig. S3B) [F(2,15)= 31.23, p<0.0001]. Our results suggested that biased recruitment of molecular effectors by SB due to different intensities may affect the future vulnerability to reinstatement.

## DISCUSSION

The present study characterized the intensity-dependent role of hippocampal TRPV1 channels in the modulation of CFC and its interaction with the ECS. First, we demonstrate that TRPV1 channels are recruited depending on the intensity of the aversive stimuli, possibly due to the progressive release of anandamide. Second, we demonstrate that even when TRPV1 channels are recruited, intensity is a key factor determining the molecular pathways engaged by TRPV1 blockers.

TRPV1 channels were already related to fear memory. For instance, blocking TRPV1 at the PFC or HPC impairs retrieval and consolidation of high intense memories (Genro et al., 2012; Terzian et al., 2014). Similarly, TRPV1 is involved in the effect of AM404, an inhibitor of the eCB transporter in high (Llorente-Berzal et al., 2015) but not moderate intensities (Bitencourt et al., 2008). Thus, AEA regulation of fear is mediated by both CB1 and TRPV1, but TRPV1 seems involved specifically in high intense memories. Equally important, AEA itself is regulated by fear memory, in this sense, the context paired with the shock enhanced AEA levels (Marsicano et al., 2002; Olango et al., 2012; Segev et al., 2018) and this release seems intensity-dependent at least after the conditioning (Morena et al., 2016). Since TRPV1 channels recruitment seems to depend on intensity as well as AEA released, and AEA effects are mediated by TRPV1 and CB1, we hypothesized that the intensity-dependent effects of TRPV1 blockers relay on the levels of AEA released due to different intensities. In accordance, we first confirmed that SB impairs freezing in moderate and high intensities but not low ones. Later, we verified that CB1 and TRPV1 are co-localized in the dHPC and that a subeffective dose of AM251, a CB1 antagonist, is able to prevent SB effect. These results suggested that the effect of TRPV1 blockers is mediated by CB1. Finally, using HPLC-MS we evaluated AEA levels immediately after retrieval in animals conditioned with different intensities and we confirm that AEA levels correlate with freezing responses. These results suggested I) that intensity-dependent release of AEA may underly the intensity-dependent recruitment of TRPV1, since this endocannabinoid acts at lower concentrations through CB1 and higher concentrations are needed to activate TRPV1; II) that CB1 mediates the impairment in freezing induced by SB.

CB1 receptors can induce fast suppression of neurotransmission (<1min) (Diana & Marty, 2004; Ohno-Shosaku et al., 2002) but also long-term forms of plasticity (Chevaleyre et al., 2006; Péterfi et al., 2012). We observed that SB treated animals did not recover the fear memory 24h after treatment, which suggest plastic modifications beyond transient inhibition of neurotransmission. In this vein, our results indicated that SB was able to enhance Arc, Zif and Trkb RNA as wells as BDNF release in the HPC, probably because of the recruitment of CB1 receptors. CB1 is able to induce long-term plasticity through pre- (Monday et al., 2020; Younts et al., 2016) and post-synaptic modifications (Maroso et al., 2016; Schweitzer, 2000). Since Arc is exclusively expressed by postsynaptic neurons (Fujimoto et al., 2004; Moga et al., 2004; Rodríguez et al., 2005), the increase of this early gene suggests some degree of postsynaptic plasticity. This is in accordance with previous reports supporting CB1 modulation of early genes in appetitive memories (Higginbotham et al., 2021). It is important to notice that despite SB impaired freezing at moderate and high intensity, the molecular alterations described were observed just in the moderate intensity group. One exception was Zif RNA levels, that were slightly enhanced in all the intensities. This is in accordance with its basal expression and its role in spatial recognition (Barbosa et al., 2013; Worley et al., 1991). In addition, we also observed an increased in Trkb-BDNF tonus induced by the treatment in the moderate intensity group. This is in agreement with several evidences pointing to a crosstalk between BDNF-TrkB and the ECS (Wu et al., 2020; Yeh et al., 2017).

Despite the mechanism involved in this divergent recruitment of molecular pathways remains unknown, a plausible explanation points to the involvement of AEA. Since moderate levels of AEA increases TrkB phosphorylation through CB1 (Diniz et al., 2019), it is possible that the AEA levels induced by the combination of moderate intensities and TRPV1 blocking lead to an enhancement of TrkB activity through CB1 receptors. Together with the higher levels of BDNF due to the treatment at this intensity, this may in turns induced the transcription of Zif, Arc and Trkb. In contrast, higher levels of AEA are available due to the combination of higher intensities and TRPV1 blocking. Since higher levels of AEA induces phosphorylation of TrkB through TRPV1 channels independent on CB1 (Diniz et al., 2019) and TRPV1 is blocked by SB, this may impair Trkb phosphorylation not increasing BDNF-Trkb tonus and, in turns, not affecting transcription.

We observed that SB is able to reduce freezing in animals conditioned with moderate and high intensities, but the molecular mechanism triggered are different dependent on the intensity of conditioning. In order to further investigate the potential consequences of these findings in fear memory, we evaluated the extinction and reinstatement of animals conditioned with moderated and high intensities and treated with SB before the retrieval. We observed that intensity did not induce differences in extinction. However, treated animals from the moderate intensity group were resistant to the reinstatement of fear memory, but this was not observed in animals conditioned with high intensities. This result suggests that biased recruitment of molecular pathways by SB due to intensity may influence the treatment effectivity in long-term basis.

Finally, since we found an increased in GDNF and immune responses were recently associated to fear memory (Chaaya et al., 2019), we evaluated the involvement of some immune markers in the CFC. First, we confirm that in the dHPC TRPV1 is not only expressed by neurons, which suggested that this channel may be express by glia cells in this structure. Later, we evaluated the levels of Cx3cr1, CX3XL1 and cytokines. We observed that Cx3cr1 RNA has a similar pattern to that observed in Arc and Trkb, which may be mediated by CB1 expressed in glia cells (Duffy et al., 2021). The levels of CX3CL1, IL-1, IL-10, Il-6 and TNFα were different between acquisition and retrieval. These findings support recent views that fear, as other injuries, induces brain neuroinflammatory responses (Chaaya et al., 2019). Unfortunately, in accordance with the invasive character of surgery and intrahippocampal administration, all the inflammatory factors evaluated were altered due to this intervention, hindering the interpretation of the results. In this sense, the inflammatory response seems to decreased 24h after the test which may be explain by the time past between the surgery and the euthanasia. However, we observed that intensity of the conditioning modifies the expression of all the inflammatory markers, which suggests some interaction between physical injury and the intensity of a psychological insult. More research is needed to better understand the involvement of inflammatory responses in fear memory and its relation with the intensity of the conditioning.

## Supporting information

Supplementary Results

## REFERENCES

Alford, G. S., & Alford, H. F. (1976). Benzodiazepine induced state-dependent learning: A correlative of abuse potential? Addictive Behaviors, 1(3), 261–267. https://doi.org/https://doi.org/10.1016/0306-4603(76)90019-8

Barbosa, F. F., Santos, J. R., Meurer, Y. S. R., Macêdo, P. T., Ferreira, L. M. S., Pontes, I. M. O., Ribeiro, A. M., & Silva, R. H. (2013). Differential Cortical c-Fos and Zif-268 Expression after Object and Spatial Memory Processing in a Standard or Episodic-Like Object Recognition Task. Frontiers in Behavioral Neuroscience, 7, 112. https://doi.org/10.3389/fnbeh.2013.00112

Bitencourt, R. M., Pamplona, F. A., & Takahashi, R. N. (2008). Facilitation of contextual fear memory extinction and anti-anxiogenic effects of AM404 and cannabidiol in conditioned rats. European Neuropsychopharmacology, 18(12), 849–859. https://doi.org/10.1016/j.euroneuro.2008.07.001

Black, N., Stockings, E., Campbell, G., Tran, L. T., Zagic, D., Hall, W. D., Farrell, M., & Degenhardt, L. (2019). Cannabinoids for the treatment of mental disorders and symptoms of mental disorders: a systematic review and meta-analysis. The Lancet. Psychiatry, 6(12), 995–1010. https://doi.org/10.1016/S2215-0366(19)30401-8

Blechert, J., Michael, T., Vriends, N., Margraf, J., & Wilhelm, F. H. (2007). Fear conditioning in posttraumatic stress disorder: evidence for delayed extinction of autonomic, experiential, and behavioural responses. Behaviour Research and Therapy, 45(9), 2019–2033. https://doi.org/10.1016/j.brat.2007.02.012

Bouton, M. E., Kenney, F. A., & Rosengard, C. (1990). State-dependent fear extinction with two benzodiazepine tranquilizers. Behavioral Neuroscience, 104(1), 44–55. https://doi.org/10.1037//0735-7044.104.1.44

Bradley, M. M., Greenwald, M. K., Petry, M. C., & Lang, P. J. (1992). Remembering Pictures: Pleasure and Arousal in Memory. Journal of Experimental Psychology: Learning, Memory, and Cognition, 18(2), 379–390. https://doi.org/10.1037/0278-7393.18.2.379

Canto-de-Souza, L., & Mattioli, R. (2016). The consolidation of inhibitory avoidance memory in mice depends on the intensity of the aversive stimulus: The involvement of the amygdala, dorsal hippocampus and medial prefrontal cortex. Neurobiology of Learning and Memory, 130, 44–51. https://doi.org/10.1016/j.nlm.2016.01.012

Caterina, M. J., Schumacher, M. A., Tominaga, M., Rosen, T. A., Levine, J. D., & Julius, D. (1997). The capsaicin receptor : a heat-activated ion channel in the pain pathway. Nature, 389(October).

Chaaya, N., Jacques, A., Belmer, A., Beecher, K., Ali, S. A., Chehrehasa, F., Battle, A. R., Johnson, L. R., & Bartlett, S. E. (2019). Contextual fear conditioning alter microglia number and morphology in the rat dorsal hippocampus. Frontiers in Cellular Neuroscience, 13(May), 1–17. https://doi.org/10.3389/fncel.2019.00214

Chadwick, V. L., Rohleder, C., Koethe, D., & Leweke, F. M. (2020). Cannabinoids and the endocannabinoid system in anxiety, depression, and dysregulation of emotion in humans. Current Opinion in Psychiatry, 33(1), 20–42. https://doi.org/10.1097/YCO.0000000000000562

Chevaleyre, V., Takahashi, K. A., & Castillo, P. E. (2006). Endocannabinoid-mediated synaptic plasticity in the CNS. Annual Review of Neuroscience, 29, 37–76. https://doi.org/10.1146/annurev.neuro.29.051605.112834

Colpaert, F. C. (1990). Amnesic trace locked into the benzodiazepine state of memory. Psychopharmacology, 102(1), 28–36. https://doi.org/10.1007/BF02245740

Cordero, M. I., Merino, J. J., & Sandi, C. (1998). Correlational relationship between shock intensity and corticosterone secretion on the establishment and subsequent expression of contextual fear conditioning. Behavioral Neuroscience, 112(4), 885–891. https://doi.org/10.1037//0735-7044.112.4.885

Cordero, M. I., & Sandi, C. (1998). A role for brain glucocorticoid receptors in contextual fear conditioning: dependence upon training intensity. Brain Research, 786(1-2), 11–17. https://doi.org/10.1016/s0006-8993(97)01420-0

Cristino, L., Petrocellis, L. D. E., & Pryce, G. (2006). Immunohistochemical localization of cannabinoid type 1 and vanilloid transient type 1 receptor in the mouse brain. Neuroscience, 139, 1405–1415. https://doi.org/10.1016/j.neuroscience.2006.02.074

Cruz-Morales, S. E., Duran-Arevalo, M., Diaz Del Guante, M. A., Quirarte, G., & Prado-Alcala, R. A. (1992). A threshold for the protective effect of over-reinforced passive avoidance against scopolamine-induced amnesia. Behavioral and Neural Biology, 57(3), 256–259. https://doi.org/10.1016/0163-1047(92)90248-3

de Oliveira, H. U., Dos Santos, R. S., Malta, I. H. S., Pinho, J. P., Almeida, A. F. S., Sorgi, C. A., Peti, A. P. F., Xavier, G. S., Reis, L. M. Dos, Faccioli, L. H., Cruz, J. D. S., Ferreira, E., & Galdino, G. (2020). Investigation of the Involvement of the Endocannabinoid System in TENS-Induced Antinociception. The Journal of Pain, 21(7-8), 820–835. https://doi.org/10.1016/j.jpain.2019.11.009

Deutsch, D. G., & Chin, S. a. (1993). Enzymatic synthesis and degradation of anandamide, a cannabinoid receptor agonist. Biochemical Pharmacology, 46(5), 791–796. https://doi.org/10.1016/0006-2952(93)90486-G

Devane, W. a, Hanus, L., Breuer, A., Pertwee, R. G., Stevenson, L. a, Griffin, G., Gibson, D., Mandelbaum, A., & Etinger, A. (1992). Isolation and Structure of a Brain Constituent That Binds to the Cannabinoid Receptor. Science, 258(10), 1946–1949. https://doi.org/10.1126/science.1470919

Diana, M. A., & Marty, A. (2004). Endocannabinoid-mediated short-term synaptic plasticity: depolarization-induced suppression of inhibition (DSI) and depolarization-induced suppression of excitation (DSE). British Journal of Pharmacology, 142(1), 9–19. https://doi.org/10.1038/sj.bjp.0705726

Diniz, C. R. A. F., Biojone, C., Joca, S. R. L., Rantamäki, T., Castrén, E., Guimarães, F. S., & Casarotto, P. C. (2019). Dual mechanism of TRKB activation by anandamide through CB1 and TRPV1 receptors. 1–21. https://doi.org/10.7717/peerj.6493

Dos Santos Corrêa, M., Vaz, B. D. S., Grisanti, G. D. V., de Paiva, J. P. Q., Tiba, P. A., & Fornari, R. V. (2019). Relationship between footshock intensity, post-training corticosterone release and contextual fear memory specificity over time. Psychoneuroendocrinology, 110, 104447. https://doi.org/10.1016/j.psyneuen.2019.104447

Duffy, S. S., Hayes, J. P., Fiore, N. T., & Moalem-Taylor, G. (2021). The cannabinoid system and microglia in health and disease. Neuropharmacology, 190, 108555. https://doi.org/10.1016/j.neuropharm.2021.108555

Eichenbaum, H., Dudchenko, P., Wood, E., Shapiro, M., & Tanila, H. (1999). The hippocampus, memory, and place cells: is it spatial memory or a memory space? Neuron, 23(2), 209–226. https://doi.org/10.1016/s0896-6273(00)80773-4

Enomoto, S., & Kato, T. A. (2021). Involvement of microglia in disturbed fear memory regulation: Possible microglial contribution to the pathophysiology of posttraumatic stress disorder. Neurochemistry International, 142, 104921. https://doi.org/10.1016/j.neuint.2020.104921

Fogaça, M. V, Aguiar, D. C., Moreira, F. A., & Guimarães, F. S. (2012). The endocannabinoid and endovanilloid systems interact in the rat prelimbic medial prefrontal cortex to control anxiety-like behavior. Neuropharmacology, 63(2), 202–210. https://doi.org/10.1016/j.neuropharm.2012.03.007

Fujimoto, T., Tanaka, H., Kumamaru, E., Okamura, K., & Miki, N. (2004). Arc interacts with microtubules/microtubule-associated protein 2 and attenuates microtubule-associated protein 2 immunoreactivity in the dendrites. Journal of Neuroscience Research, 76(1), 51–63. https://doi.org/10.1002/jnr.20056

Garín-Aguilar, M. E., Medina, A. C., Quirarte, G. L., McGaugh, J. L., & Prado-Alcalá, R. A. (2014). Intense aversive training protects memory from the amnestic effects of hippocampal inactivation. Hippocampus, 24(1), 102–112. https://doi.org/10.1002/hipo.22210

Genro, B. P., Alvares, L. D. O., & Quillfeldt, J. A. (2012). Neurobiology of Learning and Memory Role of TRPV1 in consolidation of fear memories depends on the averseness of the conditioning procedure. Neurobiology of Learning and Memory, 97(4), 355–360. https://doi.org/10.1016/j.nlm.2012.01.002

Gobira, P. H., Lima, I. V, Batista, L. A., Oliveira, A. C. De, Resstel, L. B., Wotjak, C. T., Aguiar, D. C., & Moreira, F. A. (2017). N-arachidonoyl-serotonin, a dual FAAH and TRPV1 blocker, inhibits the retrieval of contextual fear memory : Role of the cannabinoid CB1 receptor in the dorsal hippocampus. Journal of Psychopharmacology, 6, 750–756. https://doi.org/10.1177/0269881117691567

González-Franco, D. A., Ramírez-Amaya, V., Joseph-Bravo, P., Prado-Alcalá, R. A., & Quirarte, G. L. (2017). Differential Arc protein expression in dorsal and ventral striatum after moderate and intense inhibitory avoidance training. Neurobiology of Learning and Memory, 140, 17–26. https://doi.org/10.1016/j.nlm.2017.02.001

Gonzalez, S., Sagredo, O., Gómez, M., Ramos, J. A. (2002). Guía Básica sobre los Cannabinoides. Sociedad española de investigación sobre canabinoides.

Guimara, F. S., Casarotto, C., Terzian, A. L. B., & Aguiar, D. C. (2012). Opposing Roles for Cannabinoid Receptor Type-1 (CB 1) and Transient Receptor Potential Vanilloid Type-1 Channel (TRPV1) on the Modulation of Panic-Like Responses in Rats. 1, 478–486. https://doi.org/10.1038/npp.2011.207

Hartmann, A., Fassini, A., Scopinho, A., Correa, F. M. A., Guimarães, F. S., Lisboa, S. F., & Resstel, L. B. M. (2019). Role of the endocannabinoid system in the dorsal hippocampus in the cardiovascular changes and delayed anxiety-like effect induced by acute restraint stress in rats. Journal of Psychopharmacology, 33(5), 606–614. https://doi.org/10.1177/0269881119827799

Higginbotham, J. A., Wang, R., Richardson, B. D., Shiina, H., Tan, S. M., Presker, M. A., Rossi, D. J., & Fuchs, R. A. (2021). CB1 Receptor Signaling Modulates Amygdalar Plasticity during Context-Cocaine Memory Reconsolidation to Promote Subsequent Cocaine Seeking. The Journal of Neuroscience : The Official Journal of the Society for Neuroscience, 41(4), 613–629. https://doi.org/10.1523/JNEUROSCI.1390-20.2020

Hill, M. N., Bierer, L. M., Makotkine, I., Golier, J. A., Galea, S., McEwen, B. S., Hillard, C. J., & Yehuda, R. (2013). Reductions in circulating endocannabinoid levels in individuals with post-traumatic stress disorder following exposure to the World Trade Center attacks. Psychoneuroendocrinology, 38(12), 2952–2961. https://doi.org/10.1016/j.psyneuen.2013.08.004

Hitora-imamura, N., Miura, Y., Teshirogi, C., & Ikegaya, Y. (2015). Prefrontal dopamine regulates fear reinstatement through the downregulation of extinction circuits. 1–15. https://doi.org/10.7554/eLife.08274

Iglesias, L. P., Aguiar, D. C., & Moreira, F. A. (2020). TRPV1 blockers as potential new treatments for psychiatric disorders. Behavioural Pharmacology. https://doi.org/10.1097/FBP.0000000000000603

Jacob, W., Marsch, R., Marsicano, G., Lutz, B., & Wotjak, C. T. (2012). Cannabinoid CB1 receptor deficiency increases contextual fear memory under highly aversive conditions and long-term potentiation in vivo. Neurobiology of Learning and Memory, 98(1), 47–55. https://doi.org/10.1016/j.nlm.2012.04.008

Kauer, J. A., & Gibson, H. E. (2009). Hot flash : TRPV channels in the brain. Cell Press, 1(March), 215–224. https://doi.org/10.1016/j.tins.2008.12.006

Lis, S., Thome, J., Kleindienst, N., Mueller-Engelmann, M., Steil, R., Priebe, K., Schmahl, C., Hermans, D., & Bohus, M. (2020). Generalization of fear in post-traumatic stress disorder. Psychophysiology, 57(1), e13422. https://doi.org/10.1111/psyp.13422

Lisboa, S. F., Vila-Verde, C., Rosa, J., Uliana, D. L., Stern, C. A. J., Bertoglio, L. J., Resstel, L. B., & Guimaraes, F. S. (2019). Tempering aversive/traumatic memories with cannabinoids: a review of evidence from animal and human studies. Psychopharmacology, 236(1), 201–226. https://doi.org/10.1007/s00213-018-5127-x

Livak, K. J., & Schmittgen, T. D. (2001). Analysis of Relative Gene Expression Data Using Real- Time Quantitative PCR and the 2 Ϫ ◻◻ C T Method. 408, 402–408. https://doi.org/10.1006/meth.2001.1262

Llorente-Berzal, A., Terzian, A. L. B., Di Marzo, V., Micale, V., Viveros, M. P., & Wotjak, C. T. (2015). 2-AG promotes the expression of conditioned fear via cannabinoid receptor type 1 on GABAergic neurons. Psychopharmacology, 232(15), 2811–2825. https://doi.org/10.1007/s00213-015-3917-y

Lutz, B., Marsicano, G., Maldonado, R., & Hillard, C. J. (2015). The endocannabinoid system in guarding against fear, anxiety and stress. In Nature reviews. Neuroscience (Vol. 16, Issue 12, pp. 705–718). https://doi.org/10.1038/nrn4036

Mahan, A. L., & Ressler, K. J. (2012). Fear conditioning, synaptic plasticity and the amygdala : implications for posttraumatic stress disorder. Trends in Neurosciences, 35(1), 24–35. https://doi.org/10.1016/j.tins.2011.06.007

Maroso, M., Szabo, G. G., Kim, H. K., Alexander, A., Bui, A. D., Lee, S.-H., Lutz, B., & Soltesz, I. (2016). Cannabinoid Control of Learning and Memory through HCN Channels. Neuron, 89(5), 1059–1073. https://doi.org/10.1016/j.neuron.2016.01.023

Marrone, M. C., Leuti, A., Mattioli, M., Marinelli, S., Riganti, L., Lombardi, M., Murana, E., Totaro, A., Piomelli, D., Ragozzino, D., Oddi, S., Maccarrone, M., Verderio, C., & Marinelli, S. (2017). TRPV1 channels are critical brain inflammation detectors and neuropathic pain biomarkers in mice ‘ 2,4,. Nature Communications, May. https://doi.org/10.1038/ncomms15292

Marsicano, G., Wotjak, C. T., Azad, S. C., Bisogno, T., Rammes, G., Cascioll, M. G., Hermann, H., Tang, J., Hofmann, C., Zieglgänsberger, W., Di Marzo, V., & Lutz, B. (2002). The endogenous cannabinoid system controls extinction of aversive memories. Nature, 418(6897), 530–534. https://doi.org/10.1038/nature00839

Matos, M. R., Visser, E., Kramvis, I., van der Loo, R. J., Gebuis, T., Zalm, R., Rao-Ruiz, P., Mansvelder, H. D., Smit, A. B., & van den Oever, M. C. (2019). Memory strength gates the involvement of a CREB-dependent cortical fear engram in remote memory. Nature Communications, 10(1), 1–11. https://doi.org/10.1038/s41467-019-10266-1

Matsuda, L. a, Lolait, S. J., Brownstein, M. J., Young, a C., & Bonner, T. I. (1990). Structure of a cannabinoid receptor and functional expression of the cloned cDNA. Nature, 346(6284), 561–564. https://doi.org/10.1038/346561a0

Mechoulam, R., Ben-Shabat, S., Hanus, L., Ligumsky, M., Kaminski, N. E., Schatz, A. R., Gopher, A., Almog, S., Martin, B. R., Compton, D. R., Pertwee, R. G., Griffin, G., Bayewitch, M., Barg, J., & Vogel, Z. (1995). Identification of an endogenous 2-monoglyceride, present in canine gut, that binds to cannabinoid receptors. Biochemical Pharmacology, 50(1), 83–90. https://doi.org/10.1016/0006-2952(95)00109-D

Milad, M. R., Orr, S. P., Lasko, N. B., Chang, Y., Rauch, S. L., & Pitman, R. K. (2008). Presence and acquired origin of reduced recall for fear extinction in PTSD: results of a twin study. Journal of Psychiatric Research, 42(7), 515–520. https://doi.org/10.1016/j.jpsychires.2008.01.017

Moga, D. E., Calhoun, M. E., Chowdhury, A., Worley, P., Morrison, J. H., & Shapiro, M. L. (2004). Activity-regulated cytoskeletal-associated protein is localized to recently activated excitatory synapses. Neuroscience, 125(1), 7–11. https://doi.org/10.1016/j.neuroscience.2004.02.004

Monday, H. R., Bourdenx, M., Jordan, B. A., & Castillo, P. E. (2020). CB(1)-receptor-mediated inhibitory LTD triggers presynaptic remodeling via protein synthesis and ubiquitination. ELife, 9. https://doi.org/10.7554/eLife.54812

Moreira, F. A., Aguiar, D. C., Terzian, A. L. B., Guimarães, F. S., & Wotjak, C. T. (2012). Cannabinoid type 1 receptors and transient receptor potential vanilloid type 1 channels in fear and anxiety-two sides of one coin? Neuroscience, 204, 186–192. https://doi.org/10.1016/j.neuroscience.2011.08.046

Morena, M., Leitl, K. D., Vecchiarelli, H. A., Gray, J. M., Campolongo, P., & Hill, M. N. (2016). Emotional arousal state influences the ability of amygdalar endocannabinoid signaling to modulate anxiety. Neuropharmacology, 111, 59–69. https://doi.org/10.1016/j.neuropharm.2016.08.020

Morena, M., Roozendaal, B., Trezza, V., Ratano, P., Peloso, A., Hauer, D., Atsak, P., Trabace, L., Cuomo, V., McGaugh, J. L., Schelling, G., & Campolongo, P. (2014). Endogenous cannabinoid release within prefrontal-limbic pathways affects memory consolidation of emotional training. Proceedings of the National Academy of Sciences of the United States of America, 111(51), 18333–18338. https://doi.org/10.1073/pnas.1420285111

Moser, M., Rowland, D. C., & Moser, E. I. (2015). Place Cells, Grid Cells, and Memory. 1–15.

Munro, S., Thomas, K. L., & Abu-Shaar, M. (1993). Molecular characterization of a peripheral receptor for cannabinoids. Nature, 365(6441), 61–65. https://doi.org/10.1038/365061a0

Neumeister, A., Normandin, M. D., Pietrzak, R. H., Piomelli, D., Zheng, M. Q., Gujarro-Anton, A., Potenza, M. N., Bailey, C. R., Lin, S. F., Najafzadeh, S., Ropchan, J., Henry, S., Corsi-Travali, S., Carson, R. E., & Huang, Y. (2013). Elevated brain cannabinoid CB1 receptor availability in post-traumatic stress disorder: a positron emission tomography study. Molecular Psychiatry, 18(9), 1034–1040. https://doi.org/10.1038/mp.2013.61

Ohno-Shosaku, T., Tsubokawa, H., Mizushima, I., Yoneda, N., Zimmer, A., & Kano, M. (2002). Presynaptic cannabinoid sensitivity is a major determinant of depolarization-induced retrograde suppression at hippocampal synapses. The Journal of Neuroscience : The Official Journal of the Society for Neuroscience, 22(10), 3864–3872. https://doi.org/10.1523/JNEUROSCI.22-10-03864.2002

Olango, W. M., Roche, M., Ford, G. K., Harhen, B., & Finn, D. P. (2012). The endocannabinoid system in the rat dorsolateral periaqueductal grey mediates fear-conditioned analgesia and controls fear expression in the presence of nociceptive tone. British Journal of Pharmacology, 165(8), 2549–2560. https://doi.org/10.1111/j.1476-5381.2011.01478.x

Paxinos, G., & Franklin, K. (2003). The Mouse Brain in Stereotaxic Coordinates (A. Press (ed.); 2 nd).

Pedraza, L. K., Sierra, R. O., Boos, F. Z., Haubrich, J., Quillfeldt, J. A., & Alvares, L. de O. (2016). The dynamic nature of systems consolidation: Stress during learning as a switch guiding the rate of the hippocampal dependency and memory quality. Hippocampus, 26(3), 362–371. https://doi.org/10.1002/hipo.22527

Pereira, L. M., de Castro, C. M., Guerra, L. T. L., Queiroz, T. M., Marques, J. T., & Pereira, G. S. (2019). Hippocampus and Prefrontal Cortex Modulation of Contextual Fear Memory Is Dissociated by Inhibiting De Novo Transcription During Late Consolidation. Molecular Neurobiology, 56(8), 5507–5519. https://doi.org/10.1007/s12035-018-1463-4

Pertwee, R. G. (2004). Cannabinoids: Handbook of Experimental Pharmacology. In The American Journal of The Medical Sciences (Vol. 258, Issue 5). https://doi.org/10.1097/00000441-196911000-00008

Péterfi, Z., Urbán, G. M., Papp, O. I., Németh, B., Monyer, H., Szabó, G., Erdélyi, F., Mackie, K., Freund, T. F., Hájos, N., & Katona, I. (2012). Endocannabinoid-mediated long-term depression of afferent excitatory synapses in hippocampal pyramidal cells and GABAergic interneurons. The Journal of Neuroscience : The Official Journal of the Society for Neuroscience, 32(41), 14448–14463. https://doi.org/10.1523/JNEUROSCI.1676-12.2012

Ponce-Lina, R., Serafín, N., Carranza, M., Arámburo, C., Prado-Alcalá, R. A., Luna, M., & Quirarte, G. L. (2020). Differential Phosphorylation of the Glucocorticoid Receptor in Hippocampal Subregions Induced by Contextual Fear Conditioning Training. Frontiers in Behavioral Neuroscience, 14, 12. https://doi.org/10.3389/fnbeh.2020.00012

Quirarte, G. L., Cruz-Morales, S. E., Diaz del Guante, M. A., Garcia, M., & Prado-Alcalá, R. A. (1993). Protective effect of under-reinforcement of passive avoidance against scopolamine-induced amnesia. Brain Research Bulletin, 32(5), 521–524. https://doi.org/10.1016/0361-9230(93)90301-q

Rodríguez, J. J., Davies, H. A., Silva, A. T., De Souza, I. E. J., Peddie, C. J., Colyer, F. M., Lancashire, C. L., Fine, A., Errington, M. L., Bliss, T. V. P., & Stewart, M. G. (2005). Long-term potentiation in the rat dentate gyrus is associated with enhanced Arc/Arg3.1 protein expression in spines, dendrites and glia. The European Journal of Neuroscience, 21(9), 2384–2396. https://doi.org/10.1111/j.1460-9568.2005.04068.x

Ross, R. A. (2003). Anandamide and vanilloid TRPV1 receptors. British Journal of Pharmacology, 140(5), 790–801. https://doi.org/10.1038/sj.bjp.0705467

Santos, J. M., Martinez, R. C. R., & Brandão, M. L. (2006). Effects of acute and subchronic treatments with fluoxetine and desipramine on the memory of fear in moderate and high-intensity contextual conditioning. European Journal of Pharmacology, 542(1-3), 121–128. https://doi.org/10.1016/j.ejphar.2006.06.019

Schweitzer, P. (2000). Cannabinoids decrease the K(+) M-current in hippocampal CA1 neurons. The Journal of Neuroscience : The Official Journal of the Society for Neuroscience, 20(1), 51–58. https://doi.org/10.1523/JNEUROSCI.20-01-00051.2000

Segev, A., Korem, N., Mizrachi Zer-Aviv, T., Abush, H., Lange, R., Sauber, G., Hillard, C. J., & Akirav, I. (2018). Role of endocannabinoids in the hippocampus and amygdala in emotional memory and plasticity. Neuropsychopharmacology, 43(10), 2017–2027. https://doi.org/10.1038/s41386-018-0135-4

Stelt, M. Van Der, Trevisani, M., Vellani, V., Petrocellis, L. De, Moriello, A. S., Campi, B., Mcnaughton, P., Geppetti, P., & Marzo, V. Di. (2005). Anandamide acts as an intracellular messenger amplifying Ca 2 þ influx via TRPV1 channels. The Embo Journal, 24(19), 3517–3518. https://doi.org/10.1038/sj.emboj.7600839

Terzian, A. L. B., Aguiar, D. C., Guimarães, F. S., & Moreira, F. A. (2009). Modulation of anxiety-like behaviour by Transient Receptor Potential Vanilloid Type 1 (TRPV1) channels located in the dorsolateral periaqueductal gray. European Neuropsychopharmacology, 19(3), 188–195. https://doi.org/10.1016/j.euroneuro.2008.11.004

Terzian, A. L. B., Dos Reis, D. G., Guimarães, F. S., Corrêa, F. M. A., & Resstel, L. B. M. (2014). Medial prefrontal cortex transient receptor potential vanilloid type 1 (TRPV1) in the expression of contextual fear conditioning in Wistar rats. Psychopharmacology, 231(1), 149–157. https://doi.org/10.1007/s00213-013-3211-9

Tornqvist, H., & Belfrage, P. (1976). Purification hydrolyzing and Some Properties of a Monoacylglycerol- Enzyme of Rat Adipose Tissue *. The Journal of Biological Chemistry, 251(3), 813–819. http://www.ncbi.nlm.nih.gov/pubmed/1249056

Toth, A., Bocza, J., & Blumberg, P. M. (2005). Expression and distribution of vanilloid receptor 1 (TRPV1) in the adult rat brain. Molecular Brain Research, 135, 162–168. https://doi.org/10.1016/j.molbrainres.2004.12.003

Uchigashima, M., Narushima, M., Fukaya, M., Katona, I., Kano, M., & Watanabe, M. (2007). Subcellular arrangement of molecules for 2-arachidonoyl-glycerol-mediated retrograde signaling and its physiological contribution to synaptic modulation in the striatum. The Journal of Neuroscience : The Official Journal of the Society for Neuroscience, 27(14), 3663–3676. https://doi.org/10.1523/JNEUROSCI.0448-07.2007

Vouimba, R., & Maroun, M. (2011). Learning-Induced Changes in mPFC – BLA Connections After Fear Conditioning, Extinction, and Reinstatement of Fear. Neuropsychopharmacology, 2276–2285. https://doi.org/10.1038/npp.2011.115

Worley, P. F., Christy, B. A., Nakabeppu, Y., Bhat, R. V, Cole, A. J., & Baraban, J. M. (1991). Constitutive expression of zif268 in neocortex is regulated by synaptic activity. Proceedings of the National Academy of Sciences of the United States of America, 88(12), 5106–5110. https://doi.org/10.1073/pnas.88.12.5106

Wu, Y., Liu, Q., Guo, B., Ye, F., Ge, J., & Xue, L. (2020). BDNF Activates Postsynaptic TrkB Receptors to Induce Endocannabinoid Release and Inhibit Presynaptic Calcium Influx at a Calyx-Type Synapse. The Journal of Neuroscience : The Official Journal of the Society for Neuroscience, 40(42), 8070–8087. https://doi.org/10.1523/JNEUROSCI.2838-19.2020

Yeh, M. L., Selvam, R., & Levine, E. S. (2017). BDNF-induced endocannabinoid release modulates neocortical glutamatergic neurotransmission. Synapse (New York, N.Y.), 71(5). https://doi.org/10.1002/syn.21962

Younts, T. J., Monday, H. R., Dudok, B., Klein, M. E., Jordan, B. A., Katona, I., & Castillo, P. E. (2016). Presynaptic Protein Synthesis Is Required for Long-Term Plasticity of GABA Release. Neuron, 92(2), 479–492. https://doi.org/10.1016/j.neuron.2016.09.040

Zygmunt, P. M., Petersson, J., Marzo, V. Di, Julius, D., & Ho, E. D. (1999). Vanilloid receptors on sensory nerves mediate the vasodilator action of anandamide. Letters to Nature, 400(July), 6–11.

