## Supplementary Results for "Hippocampal TRPV1 channels in the modulation of contextual fear conditioning"

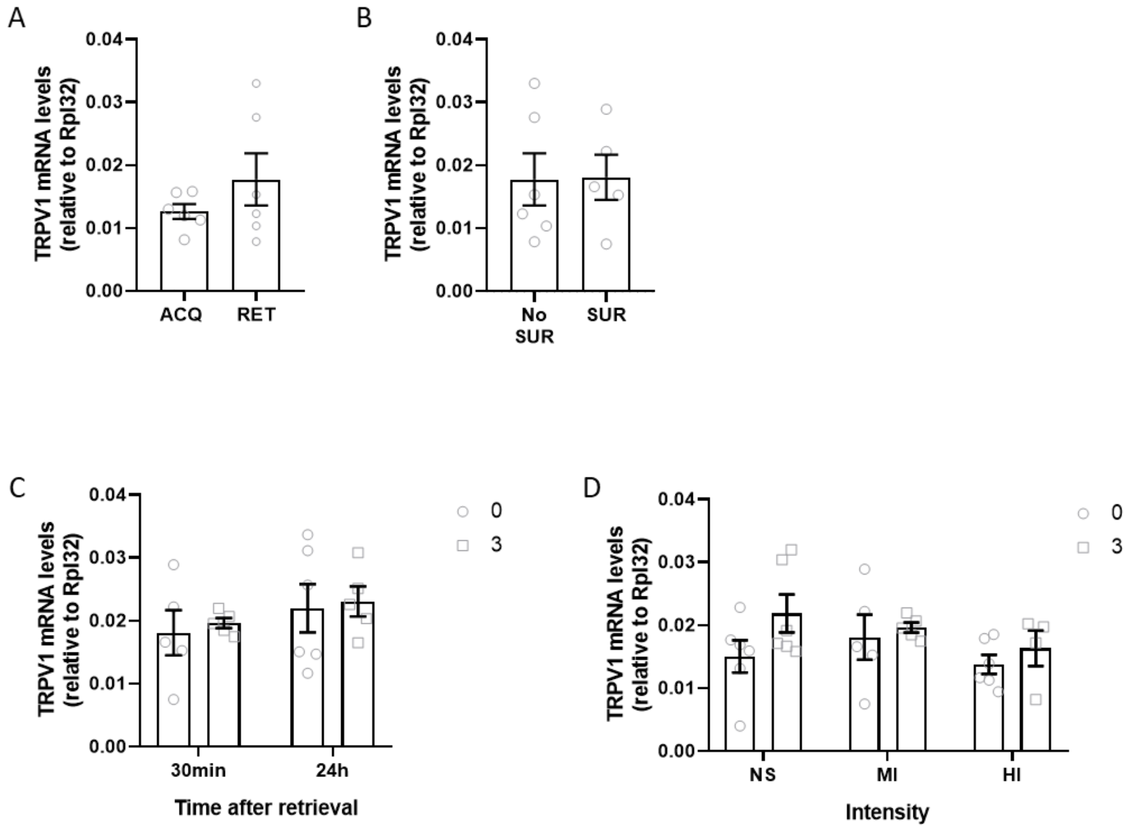


*Figure S1. RNA levels of TRPV1.* ***A)*** *RNA levels 30 min after acquisition or retrieval in animals conditioned with moderate intensity [t=1.178, df=10, p=0.266] n=6.* ***B)*** *RNA levels 30 min after retrieval in animals conditioned with moderate intensity submitted (SUR) or not to surgery (NO SUR) t-Student [t=0.06133, df=9, p=0.952], n=5-6.* ***C)*** *RNA levels 30 min or 24h after retrieval in animals conditioned with moderate intensity [Treatment: F (1,17) = 0.1842, p=0.6732. Time: F (1, 17) = 1,438, p=0.2469. Interaction: F (1,17) = 0.005418, p=0.9422] n=5-6.* ***D)*** *RNA levels 30 min after retrieval in animals conditioned with moderate intensity (MI), high intensity (HI) or no shock (NS) [Treatment: F (1, 26) = 2.978, p=0.0963. Intensity: F (2,26) = 1.254, p=0.3021. Interaction: F (2, 26) = 0.6268, p=0.5422] n=5-6.*

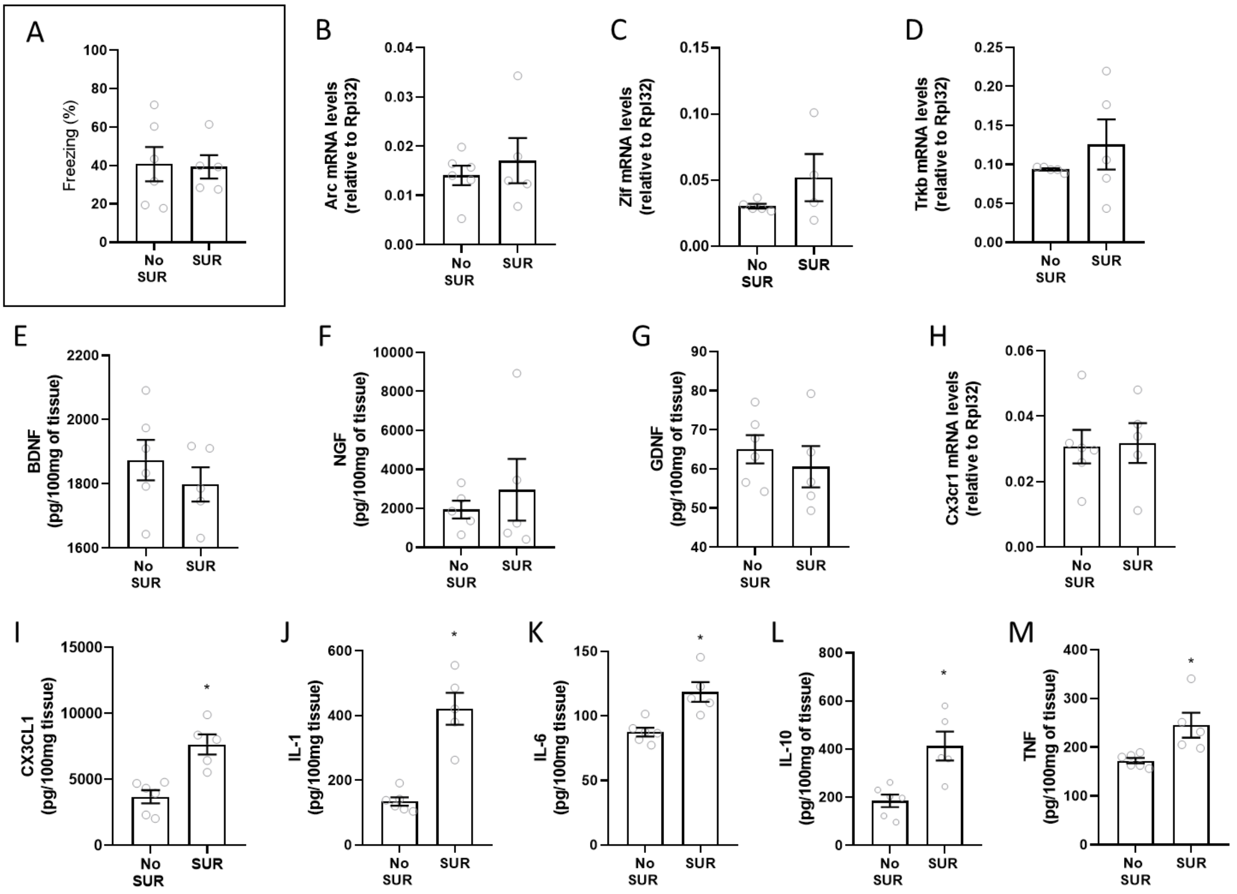


Figure S2. Early genes, neurotrophic and inflammatory factors 30min after retrieval in animals with and without surgery **A)** Freezing levels of animals with and without surgery [t=0.1246, df=9, p=0.9036] n=5-6. **B)** Arc mRNA levels [t=0.6375, df=9, p=0.5397] n=5-6. **C)** Zif mRNA levels [t=1.367, df=7, p=0.2139] n=4-5. **D)** Trkb mRNA levels [t=0.9955, df=8, p=0.3486] n=5. **E)** BDNF levels [t=0.816, df=9, p=0.3958] n=5-6. **F)** NGF levels [t=0.6176, df=8, p=0.5540] n=5. **G)** GDNF levels [t=0.7230, df=9, p=0.4881] n=5-6. **H)** Cx3cr1 mRNA levels [t=0.1373, df=9, p=0.8938] n=5-6**. I)** CX3CL1 levels [t=4.507, df=9, p=0.0015] n=5-6. **J)** IL-1β levels [t=6.117, df=9, p=0.0002] n=5-6. **K)** IL-6 levels [t=3.959 df=9, p=0.0033] n=5-6. **L)** IL-10 levels [t=3.727, df=9, p=0.0047] n=5-6. **M)** TNFα levels [t=3.052 df=9, p=0.0137] n=5-6. *p<0.05 compare to control. #p<0.05 compare to the same treatment between groups.

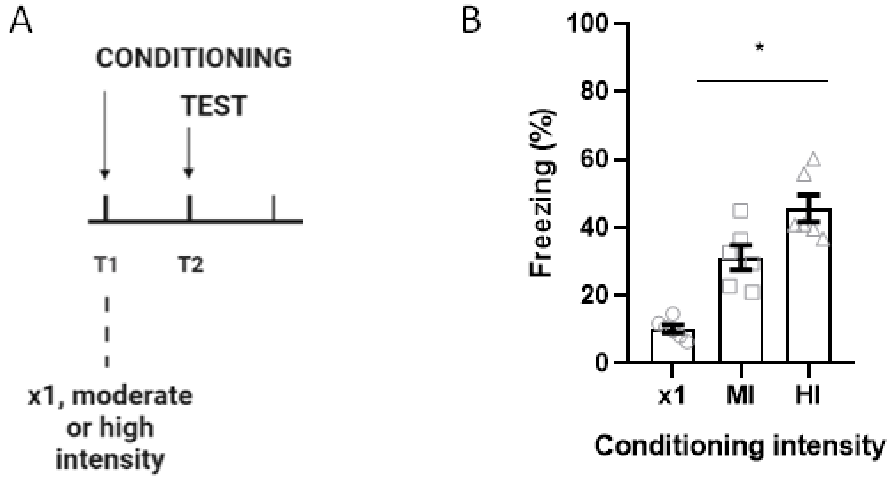


Figure S3: Freezing behaviour of animals conditioned with the reinstatement intensity (x1), moderate intensity or high intensity. **A)** Experimental design **B)** Freezing behaviour at the test [$F(2,15)= 31.23$, $p<0.0001$], n=6. *$p<0.05$ compare to control. *$p<0.05$ compare to control.
