## Supplementary Material and Methods for "Hippocampal TRPV1 channels in the modulation of contextual fear conditioning"

**Contextual Fear Conditioning**

| Name | Conditions |
|---|---|
| Low intensity | X3 shocks, 500mA, 1s |
| Moderate intensity | X3 shocks, 500mA, 2s |
| High intensity | X3 shocks, 800mA, 1s |

Table 1. Protocols used during the conditioning of CFC

### Samples obtention

The animals were euthanised by cervical dislocation. The skullcap was cut and removed. The brain was carefully removed and the hemispheres open to exposed the HPC. The HPC was dissected and kept at -80ºC until processing for HPLC-MS, PCR or ELISA.

**HPLC-MS**

As previously described by de Oliveira et al., (2020). The samples were homogenized in 500µl of MilliQ $H_2O$ by Bead ruptor (10min 30Hz). Later, 400µl were mixed with 1000µl of methanol and 10µl of the internal pattern (2AG-d5 and AEA-d4, 1000ng/ml). After a brief homogenization 500µl chloroform were added. All the volume was transferred to a solution containing 500µl of chloroform and 500µl of MilliQ water. The sample was centrifugate 10 min at 4ºC 3000rpm. The aqueous phase was collected and 500µl of chloroform was added. The sample was concentrated for 1h and resuspended in 100µl of methanol and water (7:3) and injected into the HLPC followed by the MS. The mobile phases were water and acetonitrile containing 0.1% of formic acid. The MS was operated in positive mode.

**Immunofluorescence**

The protocol was based on Fogaça et al. (Fogaça et al., 2012). Animals were prefunded with 100ml of phosphate buffered saline (PBS) and 50ml of paraformaldehyde (PFA) 4%. Coronal sections of 50 µm obtained in vibratome were kept in PFA 2% at 4ºC. The sections were washed three times of 5 min at RT with washing solution, PBS 0.01M + triton X-1000 0.3%. After that, the sections were kept in citrate buffer (pH=6) for 1h at 70ºC for antigen retrieval. After washing three times with washing solution at RT the sections were submerged in a solution of PBS 0.01M, triton X-1000 and tween 20 (1000:10:1) for 20 min at RT in order to permeabilized the sections. Later, the sections were blocking with glycine 0.1M in PBS for 20min followed by an incubation in blocking solution, BSA 5% in PBS+ 0.3% of triton X-1000. The sections were incubated in primary antibodies diluted in blocking solution for 72h: VR1 in goat (Santa Cruz, 1:30) + CB1 in rabbit (Invitrogen, 1:1000) or VR1 in goat (Santa Cruz, 1:30) + NeuN in mouse (Invitrogen, 1:500).

After 72h of incubation the sections were washed 6 times and incubated with secondary antibodies diluted in blocking solution for two hours: Alexa 594 anti-rabbit (Invitrogen, 1:1000), Alexa 488 High Cross Absorbed (Invitrogen, 1:1000) and Alexa 397 anti-mouse (Invitrogen, 1:1000). Finally, the sections were washed 6 times and mounting in gelatinize slides fixed with Flouroumnt G. The image acquisition was performed using the confocal microscopy LSM 880 Zeiss and processed by ZEN2, CB1-TRPV1 was obtained using x68, NeuN-TRPV2 using x40.

**PCR**

All the steps were performed with the samples kept in ice and in RNAse and DNAse free material unless otherwise noted.

The lyse and homogenization of the samples was performed in accordance with TRIzol® Reagent (Invitrogen) User Guide. First 500 µl of Trizol® was added to the tissue and it was homogenised with an ultrasonic homogenizer (pulse mode, 60 W), samples were kept at -20°C overnight. The day after, samples were completely defrosted and then added 100µl of chloroform (Merk Millipore). With caped tubes, the mix were homogenized by inversion during 15s. The samples were centrifugate for 15min at 12000g, the aqueous phase was transferred to a new Eppendorf containing 250 µl of isopropanol (Sigma Aldrich, molecular grade) and after a brief homogenization, the samples were centrifuged 10min at 12000 g. The supernatant was discarded by inversion and the pellet was washed in 500 µl of ethanol (Sigma Aldrich, molecular grade) and again centrifuged for 5min at 7500g. The supernatant was discarded and the RNA pellet air dried for 5min. The pellet was suspended in 20-35uL of DEPC water (LGC biotecnologia) and then incubated in a heat block for 15min at 57°C. Total RNA concentration was assessed using a nanodrop spectrophotometer (NanoDrop™ Lite, Thermo Fisher Scientific). The purity of RNA samples was determined by the ratio 260/280 absorbance (1.8 – 2 ratios were accepted). The samples were stored at -80°C until use.

M-MLV reverse transcriptase kit (Invitrogen) was used to first strand cDNA synthesis. Following manufacture's instructions, 1 µg of total RNA was used as template. All sample were diluted with DEPC water to obtained a final volume of 1000ng/µl. Later, we added 2.5 µl of MIX 1 (oligo dT, dNTP, DEPC water, 1:1:3) and the samples were kept 5 min at 65ºC. After that, the mixture was quickly chilled on ice and 4.5 µl of the MIX 2 was added (MML-V, DTT, DEPC water and M-MLV buffer, 1:2:2:3) and put in a bath at 37ºC for 50min. The reaction was inactivated by heating at 70°C for 15min. The cDNA was stored at -20ºC until use.

Quantitative PCR was performed using the iTaq™ Universal SYBR® Green Supermix (BioRad) in the CFX96 Touch™ Real Time detection system (BioRad). The PCR reaction consist on 5µl of iTaq SYBR green supermix (2x), 2µl of DNAse free water, 0,5µl of forward primer 10µM, 0,5µl of reverse primer 10µM and 2µl of cDNA 10ng/µl (10µl final volume). A mix was prepared with all reagents, except the samples, that were added in each well of PCR plate already containing 8µl of mix. The plate was sealed and centrifuged at 400g for 5 min. The PCR running conditions as well as primer sequences are described in the table 2. The running results were analysed by CFX manager and Ct values used to calculate the relative mRNA levels data Table: Oligonucleotides sequences and real time PCR running conditions.

| Target gene | Primers | Sequence 5´→ 3’ | Thermal cycling protocol |
|---|---|---|---|
| *Trpv1* | Foward | CCGGCTTTTTGGGAAGGGT | |
| | Reverse | GAGACAGGTAGGTCCATCCAC | |
| *Arc* | Foward | GTTAGCCCCTATGCCATCACC | |
| | Reverse | CTGGCCCATTCATGTGGTTCT | |
| *Erg1* (Zif/268) | Foward | TCGGCTCCTTTCCTCACTCA | 95°C/30s |
| | Reverse | CTCATAGGGTTGTTCGCTCGG | 95°C/5s, 60°C/30s – 40 X |
| *Cx3cr1* | Foward | TGCCTTCTTCCTCTTCTGGA | 65°C/5s |
| | Reverse | TAAAGGGGTTGAGGCAACAG | +0,5°C/ciclo, +0,5°C/1s – Melting |
| *Ntrk2* | Foward | CCGCTAGGATTTGGTGTACTG | |
| | Reverse | CCGGGTCAACGCTGTTAGG | |
| *Rpl32* | Foward | GCTGCCATCTGTTTTACGG | |
| | Reverse | TGACTGGTGCCTGATGAACT | |

Table 2. Oligonucleotides sequences and real time PCR running conditions.

**ELISA**

The procedure was performed as indicated by the manufacture R&D Systems kit, with some adaptations. The samples were processed in 175 µl of lysis buffer (Tris-HCl 20mM, NaCl 137mM, igepal 1%, glycerol 10%, EDTA 10mM, E-64 10mM, PMSF 1mM, pesptatin A 1µM, sodium vanadate 500mM) and homogenised with an ultrasonic homogenizer (pulse mode, 60 W). Then, they were centrifugate at 16000 rpm for 20min at 4ºC. The supernatant was collected and stored at -80°C.In order to sensibilize the plate it was added 100µl/well of capture antibody diluted 1:190 in sterile PBS and kept sealed o.n at RT. In the next day, the plate was washed three times with 300µL/well wash buffer solution (PBS + tween20 0.05%). After that, the plate was blocked with 200 µl/well of blocking solution (PBS+BSA 1%) for at least 1h at RT followed by three washes with wash buffer. The samples were diluted 1:10 in PBS+BSA 0.1% and 50 µl/well was added. The correspondent standard curve was added in this step. Sealed plate was kept at 4ºC o.n. The day after, the plate was washed 3 times and 100 µl of detection biotinylated antibody was added, the antibody was diluted 1:190 in PBS+BSA 0.1%. The plate was kept at RT for 2h. After washing three times, it was added 100 µl/well of streptavidin diluted 1:40 in PBS+BSA 0.1% and kept 30 min at RT. After washing three timed 100 µl/well of 0.3 µg/ml of OPD diluted in citrate buffer and $H_2O_2$ (5:1) was added and the plate was incubated for 30min protected from light. The reaction was interrupted with 50 µl/well of stop solution, $H_2SO_4$ 1M in distilled water. The plate was read at 490nm.
